# Structural determinants for red-shifted absorption in higher-plants Photosystem I

**DOI:** 10.1101/2025.05.05.652163

**Authors:** Stefano Capaldi, Zeno Guardini, Daniele Montepietra, Vittorio Flavio Pagliuca, Antonello Amelii, Elena Betti, Chris John, Laura Pedraza-González, Lorenzo Cupellini, Benedetta Mennucci, Diane Marie Valerie Bonnet, Antonio Chaves-Sanjuan, Luca Dall’Osto, Roberto Bassi

**Affiliations:** Dipartimento di Biotecnologie, Università di Verona, Strada Le Grazie 15, 37134 Verona, Italy; Dipartimento di Chimica e Chimica Industriale, Università di Pisa, via Giuseppe Moruzzi 13, 56124 Pisa, Italy; Accademia Nazionale dei Lincei, Palazzo Corsini, Via della Lungara 10, 00165 Rome, Italy; Anton Dorhn Experimental Marine Station, Villa Comunale, 80121 Naples, Italy; Dipartimento di Bioscienze, Università di Milano, Via Celoria 26, 20133 Milan, Italy; Fondazione Romeo e Enrica Invernizzi and Unitech NOLIMITS, Università di Milano, Via Celoria 26, 20133 Milan, Italy

**Keywords:** Far-Red, Lhca, Light-Harvesting, Low-energy absorption, Photosynthesis, Photosystem I, Red Forms

## Abstract

- Higher plants Photosystem I absorbs near-infrared light through long-wavelength chlorophylls, enriched under vegetation canopies, to enhance photon capture. Far-red absorption originates from chlorophylls pairs within the Lhca3 and Lhca4 subunits of the LHCI antenna, known as the “red cluster” composed of chlorophylls a603 and a609.
- We used reverse genetics to produce an *Arabidopsis* mutant devoid of red-shifted absorption, and we obtained high-resolution cryo-EM structures from purified PSI-LHCI complexes in both wild-type and mutant plants.
- Computed excitonic coupling values suggested a possible contribution of additional nearby pigment molecules, namely chlorophyll a615 and violaxanthin in L2 site, to far-red absorption. Therefore, we investigated the structural determinants of far-red absorption and analyzed the spectroscopic effects of these additional pigments by producing further *Arabidopsis* transgenic lines. The two experimental structures were used for quantum mechanics calculations, revealing that excitonic interactions alone cannot explain far-red absorption, while charge transfer states were needed for accurate spectral simulations.
- Our findings demonstrate that the molecular mechanisms of light-harvesting under shaded conditions rely on very precise tuning of chromophore interactions, an understanding of which is crucial for designing light-harvesting complexes with engineered absorption spectra

## INTRODUCTION

Photosynthetic organisms use solar radiation as their primary energy source to convert carbon dioxide and water into oxygen and sugars. The light reactions of oxygenic photosynthesis involve two multimeric pigment-binding protein complexes: photosystems (PS) I and II, which work in series. The initial step of light harvesting is catalyzed by PSII, which is responsible for water splitting and oxygen evolution. Meanwhile, PSI mediates the reduction of NADP^+^ to NADPH, serving as a temporary storage for reducing power (Croce & van Amerongen, 2020). Both PSs share a common structure that includes a core complex housing the reaction centers (RC) where charge separation reactions occur and an antenna system (LHC) that enhances the light-absorbing cross-section of the complexes (Pan *et al*., 2020). Despite these similarities, there are striking differences in their spectral properties, with PSI exhibiting a red-shifted absorption profile (Rivadossi *et al*., 1999). The red-most PSI absorption arises from special chlorophyll (chl) pairs that absorb at energies below that of the RC P700, referred to as “red chl forms” or simply “red forms” (RF) (Croce & Van Amerongen, 2013).

These pigments extend the absorption capacity into the near-infrared spectrum, providing advantages under canopy or in dense culture conditions where most visible wavelengths are absorbed by the upper leaf layers (Martínez-García *et al*., 2010) while infrared radiation is strongly enriched, resulting in a far-red to red ratio higher than 5 (Park & Runkle, 2017). Thus, although RF contribute only a small percentage of the total absorption (Gobets & Van Grondelle, 2001), they enjoy preferential excitation. Moreover, RF are highly effective in energy transfer and trapping, as approximately 80% of the PSI excitation transits through these pigments to reach P700 (Croce *et al*., 1998). Since these low-energy chls are highly populated, they may play crucial roles in photoprotection and/or the concentration of excitation energy (Rivadossi *et al*., 1999; Jennings *et al*., 2003; Carbonera *et al*., 2005). However, their exact physiological role(s) remain partially understood (Jennings *et al*., 2013).

RF are present in nearly all types of PSI complexes, and their occurrence is closely related to the availability of infrared radiation. In cyanobacteria (e.g., *Arthrospira platensis*), these pigments are localized within the core complex and facilitate the absorption of far-red photons, resulting in PSI fluorescence emission peaks at 720-725 nm (Karapetyan *et al*., 1997). In green algae (e.g., *Chlamydomonas reinhardtii*) and mosses (e.g., *Physcomitrium patens*), red chls are associated with the LHCI antenna system, resulting in a PSI-LHCI fluorescence emission peak wavelength ranging between 715-722 nm (Mozzo *et al*., 2010; Gorski *et al*., 2022) (Fig. **1**). Upon land colonization, environmental niches rich in far-red radiation becomes widespread under canopy, leading to a greater association of RF with LHCI, positioned distal to P700 (Croce *et al*., 1998). Indeed, the PSI-LHCI fluorescence emission of flowering plants shifts towards the red spectrum, peaking at 730 nm or even beyond (Akhtar & Lambrev, 2020).

**Figure 1.**
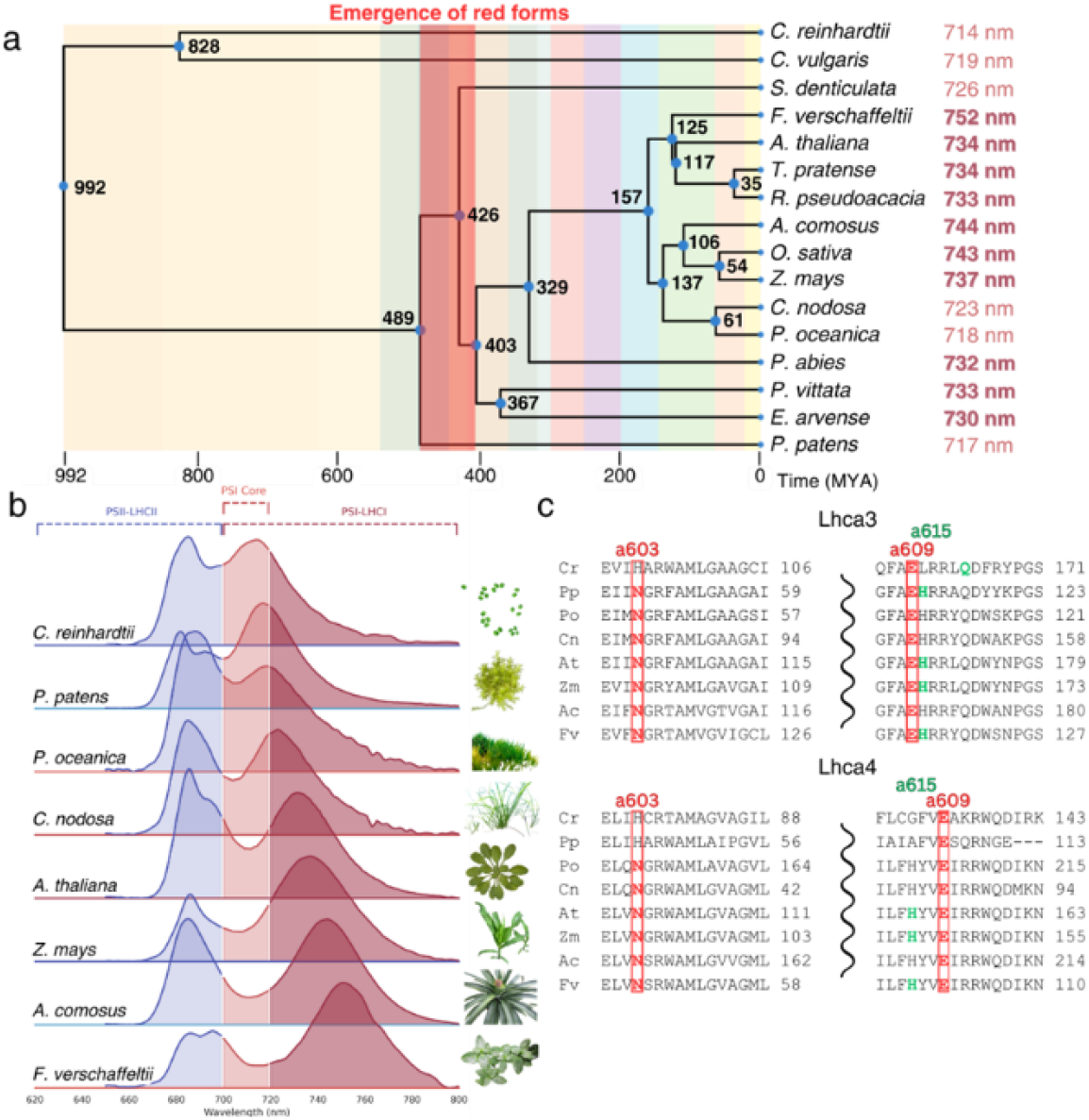
Evolution of low-absorbing spectral features in Viridiplantae. (a) Time tree of representative Viridiplantae species and their divergence times (in millions of years, MYA). The most red-shifted fluorescence emission peak of each species’ experimental spectra is reported and colored according to the color code of panel b. Note that the PSI core complex emits at ∼720 nm, which can be considered the baseline level for RF associated with antenna proteins. The emergence of RF can be traced back to approx. 489-403 MYA. (b) Low temperature (77K) fluorescence emission spectra of representative species from green algae (*C. reinhardtii*), mosses (*P. patens*), full sun land plant angiosperms: *Z. mays* (*Poaceae*) *and A. thaliana (Brassicaceae);* shade plants: *F. verschaffeltii* (*Acanthaceae*) and *A. comosus* (*Bromeliaceae*); and sea grasses (*P. oceanica* and *C. nodosa*). Shade plants thrive in far-red enriched light while seagrasses live in the absence of far-red radiation. The areas of the spectra associated with emission from PSII-LHCII, the PSI core, or the PSI-LHCI supercomplex are colored differently. (c) Multiple alignments of the Lhca4 (corresponding to Lhca8 for *C. reinhardtii* and Lhca2b for *P. patens* and Lhca3 protein sequences from *C. reinhardtii* (Cr), *P. patens* (Pp), *P. oceanica* (Po), *C. nodosa* (Cn), *A. thaliana* (At), *Z. mays* (Zm), *A. comosus* (Ac), and *F. verschaffeltii* (Fv). Residues binding red form chlorophylls (chl *a*603 and chl *a*609) are colored in red and shown in red rectangles. Coordinating residues for chl *a*615 are colored green for species with reported PDB structure.

In higher plants, LHCI binds the PSI core complex as functional heterodimers: Lhca1-Lhca4 and Lhca2-Lhca3, both of which exhibit similar spectral properties and emit far-red fluorescence at approximately 730 nm (Wientjes & Croce, 2011). *In vitro* reconstitution of recombinant Lhca complexes (rLhca) showed that the far-red shift is primarily caused by the Lhca4 and Lhca3 components of the dimers (Croce *et al*., 2002; Castelletti *et al*., 2003), and is linked to a specific sequence substitution unique to these two LHCs, where an Asn serves as the binding residue for chl *a*603 instead of a His residue, as seen in all other LHC proteins (Jansson, 1999; Morosinotto *et al*., 2003) (Fig. **1c**). Indeed, the substitution of Asn with His (*a*603-NH substitution) resulted in the complete loss of the far-red absorption and emission forms in both rLhca3 and rLhca4 complexes (Morosinotto *et al*., 2003). It was hypothesized that the presence of Asn induces strong excitonic interactions between the chl *a*603–*a*609 pair (A5 and B5 according to Kühlbrandt’s nomenclature (Kühlbrandt *et al*., 1994)) leading to a charge transfer (CT) character (Romero *et al*., 2009), and ultimately resulting in the formation of the RF (Morosinotto *et al*., 2003). Complementing *koLhca4 Arabidopsis* lines with mutant Lhca4, carrying either His or non-pigment-binding residues at the chl *a*603 ligand position (Li *et al*., 2023) resulted in a blue shift of ∼3 nm.

However, accumulating evidence suggests that the substitution of Asn *vs*. His as the ligand of chl *a*603 alone cannot account for the variability in far-red spectral properties found in nature (Li *et al*., 2023; Elias *et al*., 2024). For instance, in the Lhca2/a4/a9 subunits of *C. reinhardtii*, where the Chl *a*603 ligand is Asn, emission peaks range between 690 and 717 nm, with significantly lower absorption beyond 700 nm compared to higher plants Lhca4 (Mozzo *et al*., 2010). Moreover, the *a*603-HN (H111N) mutation, although introducing a significant red-shift in the PSII antenna complex Lhcb4, did not achieved the 43 nm shift observed in the chl *a*603-NH mutant of Lhca3 or Lhca4 (687 nm instead of ∼730 nm) (Morosinotto *et al*., 2003; Guardini *et al*., 2020; Sardar *et al*., 2024), despite the high structural similarity between the antenna complexes.

In addition to the arrangement of relevant chls, the surrounding microenvironment can be important for optimizing low-energy spectral forms. Structural models of PSI-LHCI reveal a chromophore cluster comprising three chls - chl *a*603, chl *a*609, chl *a*615 - and two xanthophylls (xan): violaxanthin in site L2 and a lutein (lut) (Qin *et al*., 2015). Notably, chl *a*615 is present only in the Lhca3 and Lhca4 subunits - the red-shifted subunits - while it is absent in all other LHC proteins, regardless of whether they serve PSI or PSII. This observation raised the question of its potential role in forming excitonic interactions (Melkozernov & Blankenship, 2003; Wientjes *et al*., 2012).

In this study, we investigated the structural and biophysical determinants of the red chl forms in *Arabidopsis thaliana* PSI-LHCI by combining *in vivo* site-directed mutagenesis and single-particle cryogenic electron microscopy (cryo-EM). We obtained two high-resolution structures of the PSI-LHCI supercomplexes from *Arabidopsis thaliana* wild-type (WT) and the *a*603-NH mutant genotype, which lacks RF. We then computed the excitonic interactions within PSI-LHCI pigments, focusing on the interaction between the L2 xan and the chl *a*603– *a*609 pair in forming low-lying energy spectral states.

## MATERIALS AND METHODS

### Plant materials

The *Arabidopsis thaliana koLhca3 koLhca4* mutant was generated by crossbreeding *koLhca3* and *koLhca4* NASC insertional lines, following the methodology outlined by (Bressan *et al*., 2016). Complementation of the *koLhca3 koLhca4* mutant was achieved through *Agrobacterium tumefaciens*-mediated transformation, as described by (Zhang *et al*., 2006), resulting in *a603-NH, a615-HA, a615-HI* and *A3WT-A4WT* lines.

### Purification and characterization of PSI-LHCI samples

To purify PSI-LHCI supercomplexes, *Arabidopsis thaliana* plants were grown for approximately 6 weeks in a phytotron (150 μmol photons m^−2^ s^−1^, 23°C, 70% relative humidity, 8/16 h of day/night). Prior to the isolation procedure, *Arabidopsis* leaves were dark-adapted for 60 minutes at 4°C. Unstacked thylakoid membranes were prepared as described in (Bassi *et al*., 1985). Thylakoid membranes were resuspended to a chl concentration of 1 mg/mL in 10 mM HEPES pH 7.5 and solubilized by adding an equal volume of 2% dodecyl-β-D-maltoside (β-DM). The samples were vortexed for 30 seconds and incubated on ice for 10 minutes, and the insoluble material was removed by centrifugation at 20,000 xg for 10 minutes at 4°C. The supernatants were fractionated by sucrose gradient ultracentrifugation at 284,000 xg at 4°C for 18 hours (Beckman SW40 Ti rotor) or at 141,000 xg at 4°C for 30 hours (Beckman SW28 Ti) (Fig. **S1**). Bands containing PSI-LHCI were harvested using a Hamilton syringe. For cryo-EM preparations, PSI-LHCI samples were concentrated to a final volume of ∼800 μL, and sucrose was removed by dialysis overnight against a solution containing 10 mM HEPES pH 7.5 and 0.05% β-DM. Finally, the samples were concentrated to a chl concentration of ∼1,5-2 mg/mL.

### Purification PSI-core and LHCI

PSI-core complex and LHCI were purified from WT and mutant lines as described by (Croce *et al*., 1998; Wientjes & Croce, 2011) with some modifications. Briefly, PSI-LHCI from sucrose gradient ultracentrifugation were diluted in 10 volumes of 5 mM tricine (pH 7.8) and centrifuged for 3 h at 411,000 xg, using a T-865.1 rotor, Sorvall. Pellets were resuspended in a buffer containing 5 mM Tricine (pH 7.8) and 0.05% β-DM. The chl concentration was adjusted to 0.3 mg/mL and the samples were solubilized by adding 1% β-DM and 0.5% Zwittergent-16. The samples were kept on ice with gentle agitation for 25 min, then rapidly frozen in liquid nitrogen and slowly thawed to enhance the yield in detached LHCI. Solubilized complexes were fractionated by sucrose gradient ultracentrifugation at 485,000 xg at 4°C for 6 hours, using a Beckman SW60 Ti rotor.

### Spectroscopy and pigment analysis

Absorption spectra were recorded at room temperature (RT, 22°C) using an SLM-Aminco DW-2000 spectrophotometer in a buffer containing 10 mM HEPES pH 7.5 and 0.05% β-DM for native complexes or 80% acetone buffered with Na_2_CO_3_ for pigment extracts.

Emission and excitation fluorescence spectra were recorded at cryogenic temperatures (77 K) using a Jobin–Yvon Fluoromax-3 spectrofluorometer. Samples were diluted in a buffer containing 50% w/v glycerol, 10 mM HEPES, pH 7.5, 0.05% β-DM, and excited at 440 nm.

Emission spectra of intact leaves were recorded at 77 K using an Ocean Insight SR-6NVN500-50 spectrofluorometer.

CD spectra were recorded at 4°C on a Jasco J1500 spectropolarimeter.

The pigment composition of LHCI complexes was assessed from the deconvolution of acetonic spectra as described in (Chazaux *et al*., 2022). HPLC pigment separation and quantification was performed according to (Gilmore & Yamamoto, 1991) by using an Agilent 1260 Infinity II HPLC system.

### Cryo-EM sample preparation and data acquisition

The PSI supercomplex at a chl concentration of ∼1.5-2 mg/mL was vitrified with a Mark IV Vitrobot (Thermo Fisher Scientific). 3 μL of the sample were applied to a Quantifoil R 0.6/1Cu 300-mesh grid previously glow-discharged at 30 mA for 30 seconds in a GloQube (Quorum Technologies). Immediately after sample application, the grids were blotted in a chamber at 4 °C and 100% humidity and then plunge-frozen into liquid ethane.

Vitrified grids were transferred to a Talos Arctica (Thermo Fisher Scientific) operated at 200 kV and equipped with a Falcon 3 direct electron detector (Thermo Fisher Scientific). 2716 and 2601 movies were acquired, for WT and *a603-NH* mutant, respectively, at a nominal magnification of 120’000x, corresponding to a pixel size of 0.889 Å/pixel, in electron counting mode, with a nominal defocus range of -0.8 to -2.4 μm and with a total dose of 40 e^−^ /Å^2^ equally distributed on 40 frames. The cryo-EM experiments were conducted at the NoLimits Center of the University of Milan.

### Cryo-EM data processing and image reconstruction

All image processing and reconstruction steps were performed using CryoSPARC v4.4 (Punjani *et al*., 2017). The experimental workflows are outlined in Fig. **S2** and **S3**. After patch motion correction and CTF estimation, 2313 and 2257 manually curated micrographs were used for initial particle picking for WT and *a603-NH* mutant, respectively.

For AtPSI-WT, after a first round of reference-free autopicking and 2D classification, the best classes were used for template-based autopicking, resulting in an initial number of 575,271 particles. Particles were extracted with a box size of 448 px, binned 2X2, and subjected to several rounds of 2D classification. The best 2D classes (282,376 particles) were used for ab initio reconstruction (3 classes) and heterogeneous refinement, resulting in an initial map at 3.74 Å resolution. After 3D classification and local and global CTF refinement, the best particles were re-extracted at full size and used for a final round of non-uniform refinement (Punjani *et al*., 2020), obtaining a final density map at 3.13 Å resolution.

For AtPSI-*a*603-NH, 531,976 particles generated from a first round of reference-free autopicking were subjected to 2D classification, and the best classes were used to train the neural network in Topaz (Bepler *et al*., 2019) on a subset of 157 micrographs. The trained model was then used to pick the entire dataset, resulting in a total number of 115,589 particles. These particles were extracted with a box size of 448 px, binned 2X2, and after several rounds of 2D classification, the best particles were selected for ab initio reconstruction (3 classes) and heterogeneous refinement, resulting in an initial 4.19Å resolution map. After two rounds of 3D classification, particles corresponding to the best 3D class were re-extracted without downscaling and, following local and global CTF refinement, further subjected to homogeneous and non-uniform refinement, yielding a final map with a resolution of 3.29 Å.

Before model building, the maps were sharpened with a global B factor of 83.7 Å^2^ and 91.8 Å^2^ for WT and *a603-NH* mutant, respectively. The resolution was estimated by the “gold-standard” Fourier Shell Correlation (FSC=0.143) criterion. Local resolution estimation was performed as implemented in CryoSPARC on the unsharpened map. The final EM maps (colored according to the local resolution) and the FSC curves are shown in Fig. **S2** and **S3** (panels D and E).

### Model building and refinement

The high-resolution structure of the *P. sativum* PSI-LHCI complex (PDB code 7DKZ) was used as the starting model. After initial docking of the model in the map with USCF ChimeraX (Goddard *et al*., 2018), each protein chain was independently rigid-body fitted and refined with Phenix-refine (Adams *et al*., 2010). The amino acid sequences of the different subunits were manually mutated to match the sequences of *A. thaliana* using COOT (Emsley & Cowtan, 2004), and the resulting model underwent several rounds of real-space refinement in Phenix and manual rebuilding in COOT. Ligands and water molecules were modelled when unambiguously identified in the density map and refined to a reasonable B factor. Ligand restrains for refinement were generated with eLBOW (Moriarty *et al*., 2009). The stereochemical quality of the final model was assessed with MolProbity (Chen *et al*., 2010). Data collection and model refinement statistics are summarized in Table S3. High-resolution figures were prepared with ChimeraX and PyMOL (The PyMOL Molecular Graphics System, Version 2.0 Schrödinger, LLC).

### Energy transfer and excitonic coupling calculations

We used a point-dipole approximation (PDA) to compute the excitonic couplings, as described by (van Amerongen & van Grondelle, 2001; Müh *et al*., 2010; Liguori *et al*., 2015; Sen *et al*., 2021). The chl point dipoles were positioned at the geometric centers of the four nitrogen atoms of the chlorin ring (Friedl *et al*., 2022)

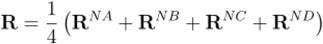

The chl Q_Y_ and Q_X_ transition dipole moments were aligned parallel to the nitrogen ND-NB and NC-NA axis, respectively, as in (van Amerongen & van Grondelle, 2001; Georgakopoulou *et al*., 2007; Liguori *et al*., 2015). The carotenoids (car) point dipole was placed on the C15 atom, with transition dipole moments (transition S2 ← S0) oriented parallel to the central part of the polyene chain (atoms C11-C33), as modelled in previous studies (Georgakopoulou *et al*., 2007; Liguori *et al*., 2015). The Q_Y_ transition dipole moments were used for couplings between chls (chl *a*–chl *a*, chl *a*–chl *b*, and chl *b*–chl *b*), while couplings between chls and cars were computed with the Q_X_ transition dipole moments for chls (Croce *et al*., 2001; Polívka & Frank, 2010).

The excitonic couplings between two pigments i and j, in the PDA were calculated using the following formula (cm^−1^):

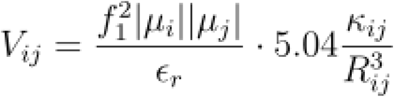

where f_1_ is the local field correction factor, |*μ*_*i*_| and |*μ*_*j*_|are the module of the transition dipole moment of pigments i and j, ε_r_ is the relative dielectric constant, here equal to 2.4 (Gobets & Van Grondelle, 2001; Liguori *et al*., 2015) *R*_*ij*_ is the module of the distance between the center of the dipole moment vector and *k*_*ij*_ is the orientation factor between chl i and j

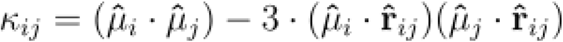

with 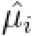 and 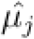 are the normalized transition dipole moment vector and 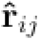 is the normalized distance vector. Dipole moment values were taken as 4 D, 3.4 D, and 4.5 D for chl *a*, chl *b*, and cars, respectively (van Amerongen & van Grondelle, 2001; Georgakopoulou *et al*., 2007; Liguori *et al*., 2015) In the case of ε_r_ = 2.4, (f_1_^2^μ^2^)/ε_r_ was calculated to be 17.6 D^2^ for chl *a* (van Amerongen & van Grondelle, 2001). The calculations were run via homebuilt codes using Python 3.8.

### Excited state calculations

We performed QM/MM optimizations and polarizable QM/MM (Bondanza *et al*., 2020) excited-state calculations of the Chls in the Lhca4 structure from the present work. Before QM/MM calculations, the structure was refined in a pure MM protocol as detailed in the Supplementary Information. All chls were optimized independently, except for *a*603-*a*609 which were optimized together. Excited-state calculations were performed for all chls in Lhca4 and for the *a*603-*a*609 dimer; control calculations were performed by also including L2 Vio with chls *a*603 and *a*609. We employed a diabatization procedure to extract CT energies and couplings from the dimer calculations (Nottoli *et al*., 2018). Finally, we built an exciton model considering all Q_Y_ states of chls and the CT states within the *a*603-*a*609 dimer, which we used to simulate absorption spectra of Lhca4 (Sláma *et al*., 2023). Detailed computational methods are reported in the SI.

### Statistics

Statistical analyses were performed in OriginPro using One-way analysis of variance (ANOVA), means were separated with Tukey’s post hoc test at a significant level of P < 0.05 (see figure legends for details). Error bars represent the standard deviation.

## RESULTS

The *koLhca3 koLhca4 Arabidopsis thaliana* genotype was complemented with mutant isoforms of Lhca3 and Lhca4, in which the Asn responsible for chl *a*603 binding was replaced with a His (*a*603-NH mutant) in both antenna subunits (Asn99 in AtLhca4 and Asn103 in AtLhca3). We observed a shift in the 77K fluorescence emission spectrum of the leaves from 736 to 724 nm (Fig. **S4**).

PSI-LHCI supercomplexes from both WT and *a*603-NH mutant lines were purified by sucrose gradient ultracentrifugation (Fig. **S1**). RT absorption spectra confirmed the loss of far-red spectral forms in the *a*603-NH mutant complex, which peaked at 704 nm and extended to 750 nm. This was accompanied by an increased absorption ranging from 650 to 700 nm, peaking at 679 nm (Fig. **2a**). Low temperature (77K) emission spectra revealed a 13 nm blue shift in the *a*603-NH mutant PSI supercomplexes compared to WT, with emission peaks at 721 nm (Fig. **2b**). Emissions around 720 nm can be attributed to the PSI core complex (Bassi & Simpson, 1987), leading us to conclude that the *a*603-NH mutation effectively abolished Lhca-associated RF *in vivo*, consistent with previous reports (Morosinotto *et al*., 2003; Li *et al*., 2023).

**Figure 2.**
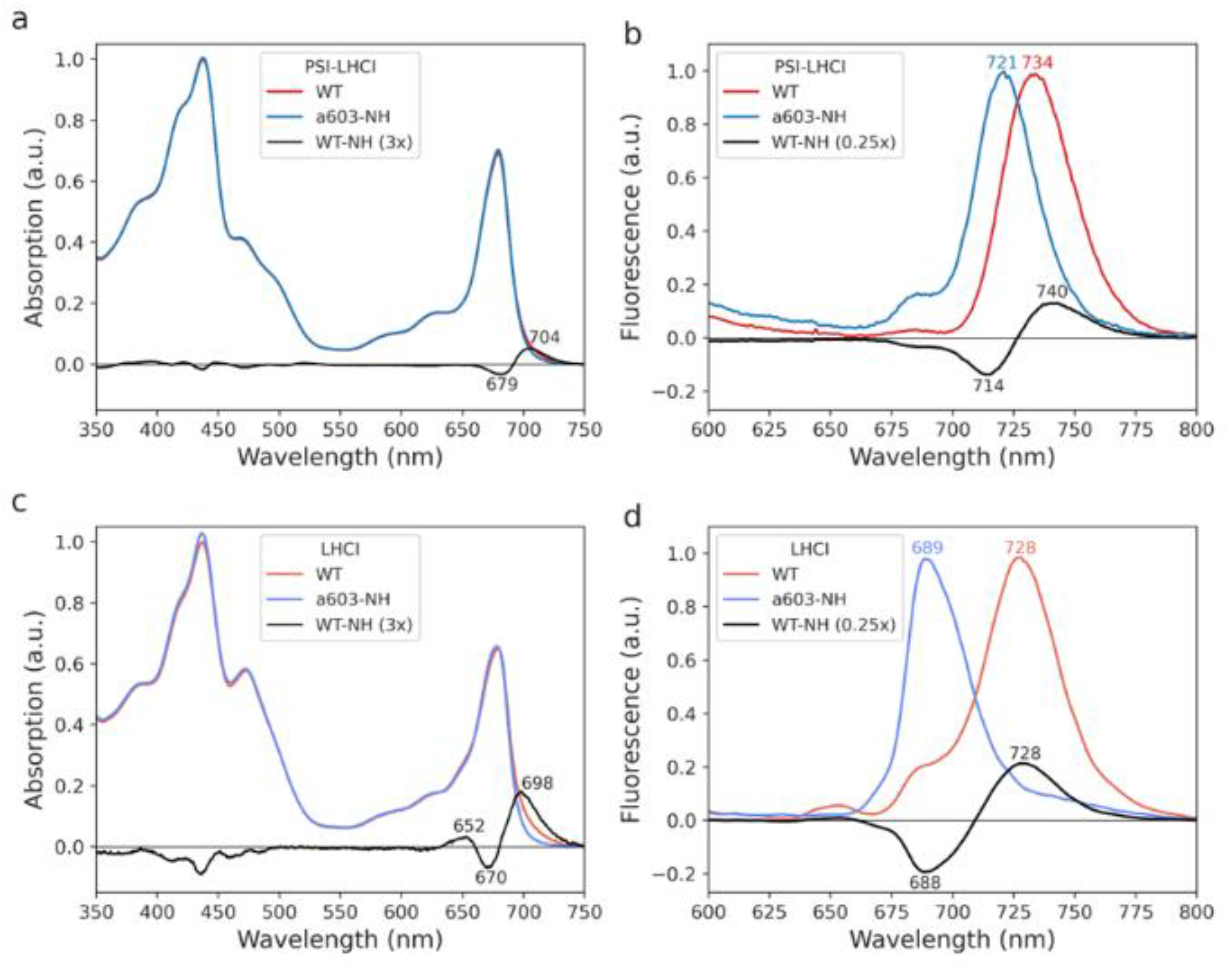
Spectral analysis of PSI-LHCI and LHCI from *A. thaliana* WT and *a603-NH*. (a, c) RT absorption and (b, d) 77K fluorescence emission spectra of PSI-LHCI (a, b) and LHCI (c, d) from *A. thaliana* WT and *a*603-NH. The WT minus NH difference spectra are shown as black lines in the corresponding plots. The amplitude of the absorption difference spectra was magnified by a factor of 3, while the fluorescence difference spectra were multiplied by a factor of 0.25 in order to plot them on the same axis. Key wavelengths are indicated in nm above the respective peaks. Experiments were repeated independently twice, with similar results.

To assess the impact of the *a*603-NH mutation on the spectral properties of LHCI complexes compared to PSI-LHCI supercomplexes, we purified LHCI heterodimers (Lhca1-Lhca4 and Lhca2-Lhca3). LHCI dimers isolated from WT plants exhibited a 77K emission maximum at 728 nm, consistent with (Wientjes & Croce, 2011). In contrast, LHCI dimers from the *a*603-NH line showed an emission peak at 689 nm, highlighting a significant blue shift of approximately 39 nm (Fig. **2d**), which aligns with observations made *in vitro* (Morosinotto *et al*., 2003). Similarly, RT absorption spectra of isolated LHCI showed the loss of a broad absorption tail in the *a*603-NH mutant, covering the range from 680 to 730 nm and peaking at 698 nm (Fig. **2c**). Conversely, the PSI core purified from both WT and *a*603-NH lines exhibited unchanged optical properties, maintaining a peak at 720 nm (Fig. **S5**).

### Structures of PSI-LHCI from WT and a603-NH plants

To investigate the structural determinants underlying the blue-shifted absorption/emission caused by the *a*603-NH mutation, we determined the cryo-EM structures of the *A. thaliana* PSI-LHCI WT and *a*603-NH supercomplexes. The final reconstructions, at 3.13 Å and 3.29 Å resolution, respectively, revealed well-defined density maps for both the PSI core and the LHC subunits (Fig. **S6**), allowing the construction of accurate models for the two complexes.

The structure of the individual subunits and the positioning of the chromophores within the *At*PSI-LHCI WT supercomplex were very similar to those observed in other land plants (Qin *et al*., 2015; Mazor *et al*., 2017; Huang *et al*., 2021; Iwai *et al*., 2024; Nelson, 2024) (Fig. **S7-S12**, Table **S1**). In both *At*PSI-LHCI WT and *a*603-NH structures, the primary difference in pigment composition between the different Lhcas was found in the chl a/b ratio (Fig. **S13**). Specifically, Lhca1/a2/a3 each bound 14 chl, while Lhca4 bound 15 chls.

Chl a615 was coordinated by His168 and His151 from helix C in Lhca3 and Lhca4, respectively. These pigments showed clear density maps in both structures, allowing for accurate modelling of the chls (including the chlorin rings and parts of the phytol tails) and the xan molecule (Fig. **S14**). An additional xan molecule (lut) was located close to chl *a*615 in Lhca4 only. Since this lut was absent in the red-emitting Lhca3, it was not considered a potential component of the red-emitting cluster.

In the *a*603-NH mutant structure, the bulkier His side chain at position 103 (Lhca3) and 99 (Lhca4) caused a ∼0.7 Å displacement of the chl *a*603 chlorin ring (Fig. **3**, **S14, S15, S16, S17**), clearly visible in the corresponding density maps (Fig. **S14**). The chlorin ring and part of the phytol tail of chl *a*603 were well-defined in both the WT and *a*603-NH density maps, allowing the reliable assignment of the position of chl *a*603 in both structures (Fig. **S14b, c**). Surprisingly, the distance between the chlorin ring centers of chl *a*603 and chl *a*609 remained essentially unchanged across both WT and mutant proteins (9.3 Å in WT *vs*. 9.1 Å in the *a*603-NH for Lhca3, and 9.3 Å *vs*. 9.2 Å for Lhca4). In comparison, the distance between the centers of the closest pyrrolic rings of the two chls (ring C) increased from 4.2 to 4.4 Å in Lhca3 and from 4.3 to 4.6 Å in Lhca4 (Fig. **S15**).

**Figure 3.**
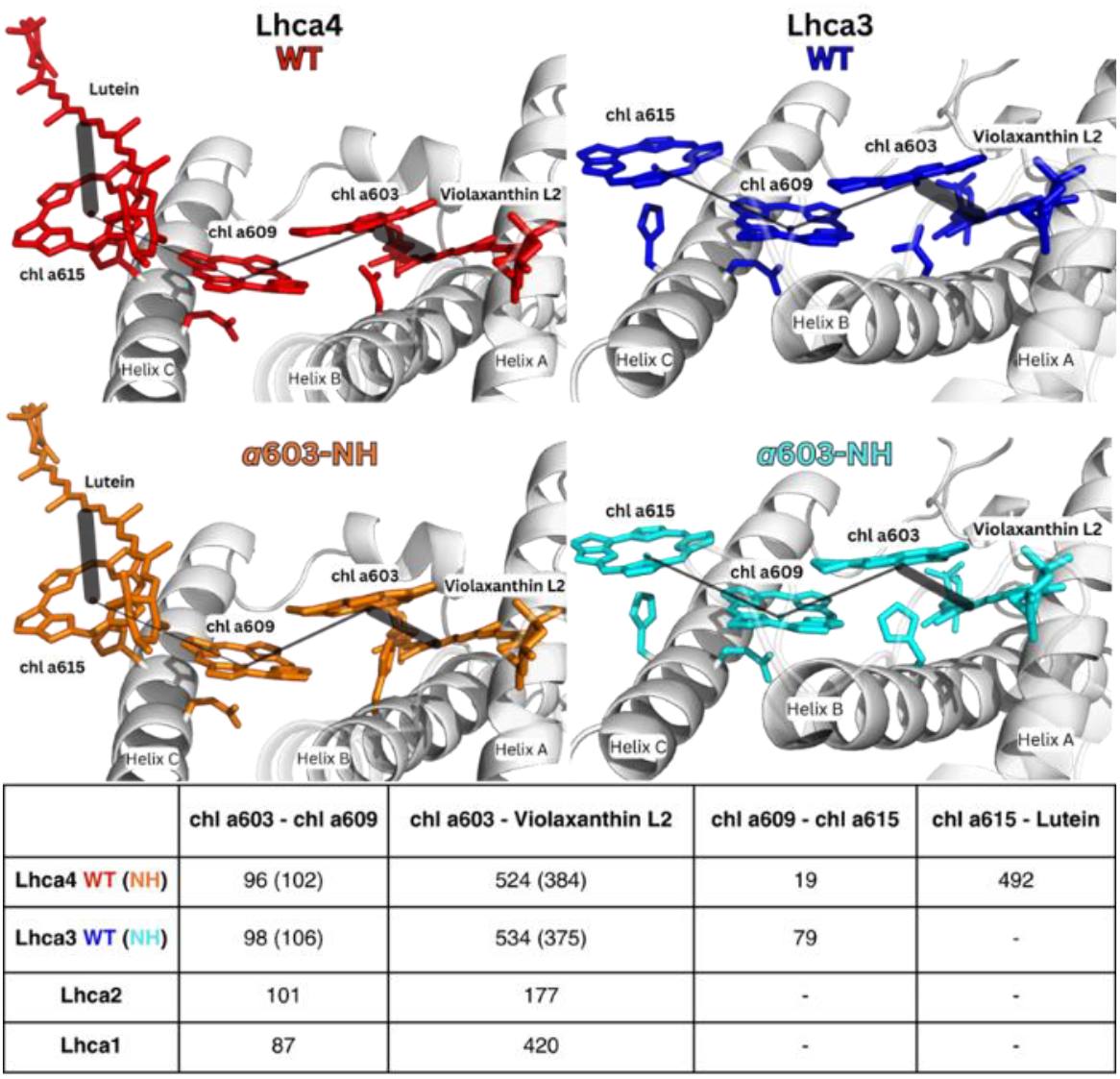
Structural superposition and excitonic coupling analysis of Lhca4 and Lhca3 WT and *a603-NH*. (*upper part*) Structures of WT and *a*603-NH red clusters of Lhca4 and Lhca3 from *A. thaliana* PSI-LHCI. WT structures are colored in red and orange, while *a603-NH* mutant structures are colored in blue and cyan. Lines are drawn between the pigments to highlight the excitonic couplings reported in the table (*lower part*, values in cm^−1^). The thickness of the line is proportional to the EC value.

At the same time, the distance between chl a603 and violaxanthin in L2 increased from 5.2 Å in WT to 5.9 Å in *a*603-NH, and from 5.3 Å to 6.0 Å in both Lhca3 and Lhca4, representing a change of about 12%.

We then computed excitonic coupling (EC) values, which take into account the relative spatial orientation between each pair of pigments in the cluster. The EC between chl *a*603 and Vio decreased from 534 cm^−1^ and 524 cm^−1^ for Lhca3 and Lhca4 in the WT to 375 cm^−1^ and 384 cm^−1^ in the *a*603-NH mutant, representing a change of 27-29% (Fig. **3**, **S18**). In contrast, the introduction of the *a*603-NH mutation resulted in only slight alterations of the EC values between the chl *a*603 and chl *a*609 pair (6-8 cm^−1^). Furthermore, while the EC values between the chl pairs were in the limited range of 87 cm^−1^ and 101 cm^−1^ for all Lhca, the EC values for the WT chl *a*603–Vio L2 coupling were significantly higher for Lhca3 and Lhca4 compared to Lhca1 and Lhca2, which do not show red-shifted absorption. Notably, the *a*603-NH mutation decreased the ECs of Lhca3 and Lhca4 to values comparable to those of the ‘non-red’ subunits. Collectively, these data suggest that the xan ligand might play a role in tuning the absorption properties of Lhca3 and Lhca4 toward low energy levels.

### Effect of Xanthophyll occupancy of site L2 on low-energy absorption

To investigate the role of the xan at the L2 site of Lhca3 and Lhca4 in modulating RF-inducing interactions, we compared the fluorescence excitation spectra of WT and *a*603-NH PSI-LHCI, highlighting the contribution of different wavelengths and associated chemical species to their far-red fluorescence emissions. In the 455-505 nm region, with the major contribution from cars (Ashenafi *et al*., 2023), the *a*603-NH mutant showed a systematically lower signal compared to WT (Fig. **4a**), implying a change in the efficiency of energy transfer from xan to far-red chl.

**Figure 4.**
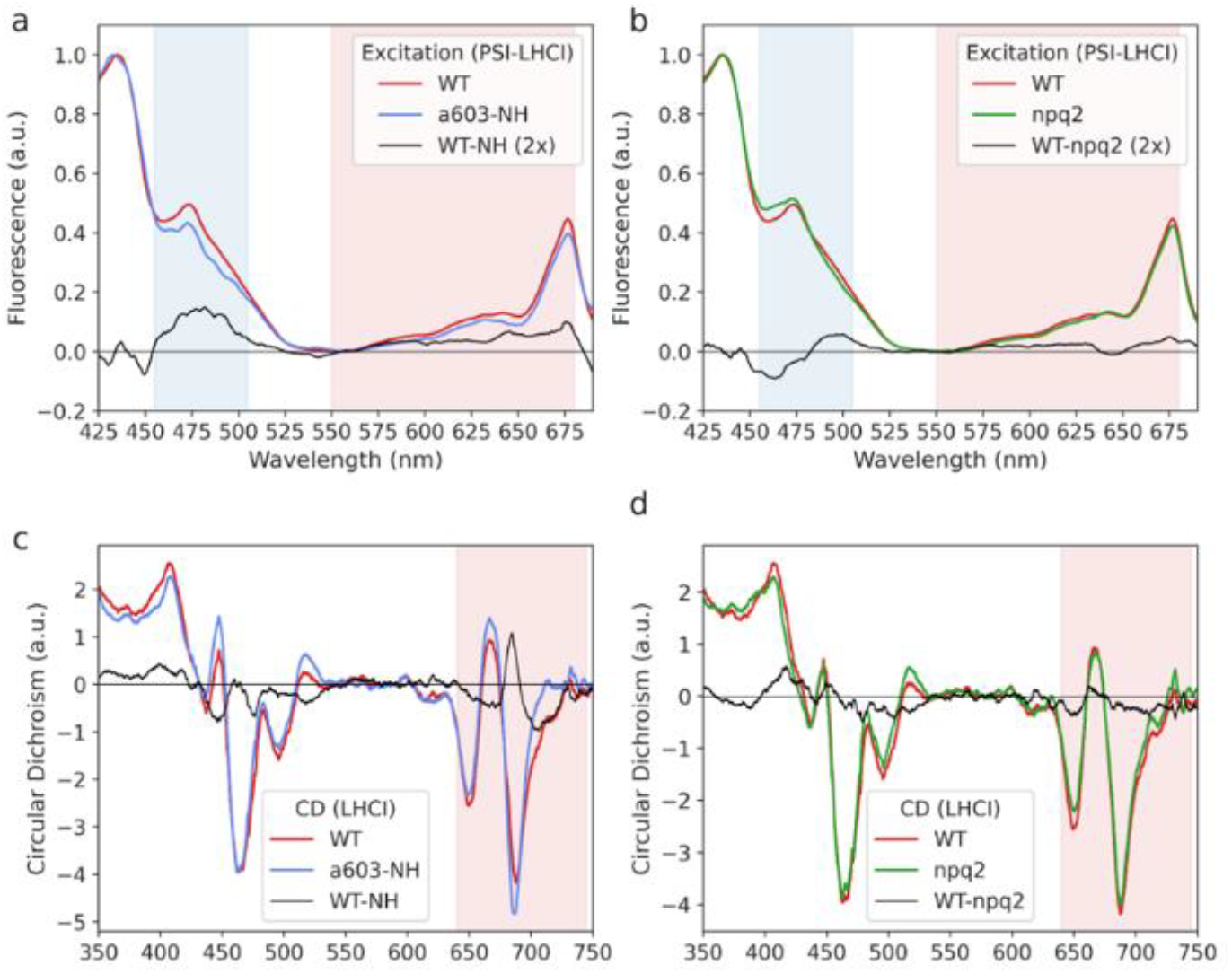
Spectral analysis of PSI-LHCI and isolated LHCI from A. thaliana WT, *a*603-NH, and *npq2* plants. Low-temperature (77K) fluorescence excitation spectra of PSI-LHCI from A. thaliana WT, (a) *a*603-NH, (b) and npq2 plants. Far-red fluorescence emission (at 760 nm for WT and *npq2*, and 740 nm for the *a*603-NH mutant) was followed by exciting the samples from 425 to 690 nm. Spectra were normalized to the peak value. Shaded areas indicate spectral regions where carotenoids and chls absorption is dominant and are colored in blue and red, respectively. Key wavelengths corresponding to absorption/emission peaks are indicated in nm above the respective peaks. c-d) CD (4°C) spectra of purified LHCI complexes from WT, *a*603-NH and *npq2* mutants. Spectra are normalized to the same absorption in the Q_Y_ region. The difference spectra are shown as black lines in the corresponding plots. The values of the fluorescence excitation difference spectra have been magnified by a factor of 2 for better visualization. Experiments were repeated independently twice, with similar results.

The difference between absorption and excitation (Abs-Exc) plots in Fig. **S19a** provides insight into the extent of energy absorbed and then transferred to chl *a* at each wavelength. In the 475-505 nm region, the Abs *minus* Exc values for the a603-NH mutant were larger than those of the WT, suggesting stronger energy dissipation (i.e., lower conversion to chl *a* excited state) and a reduced overall contribution of cars to far-red fluorescence.

Changes were also observed in the 560-680 range, which is associated with contributions from chl (Ashenafi *et al*., 2023). The *a*603-NH mutant also showed a lower signal compared to the WT. Specifically, the highest peak and the area under the curve showed a decrease of 4% and an 8%, respectively, in the *a*603-NH *vs*. the WT.

We proceed to investigate the role of xan in promoting RF. Since the removal of xan in site L2 is not feasible due its essential role in protein folding and stability (Dall’Osto *et al*., 2013), we analyzed the effect of altering L2 occupancy. To this aim, we isolated the PSI-LHCI supercomplex from the *npq2* genotype, which has zeaxanthin (zea) replacing vio (Ballottari *et al*., 2014) owing to the inactivation of zeaxanthin epoxidase (Niyogi *et al*., 1998) (Fig. **S15**). The low-temperature fluorescence emission spectra revealed a ∼2 nm blue shift in the *npq2* PSI-LHCI compared to the WT supercomplex (Fig. **S20b**).

We then recorded fluorescence excitation spectra to assess the different contributions to fluorescence emission within the wavelength range of 450 to 520 nm for both the *npq2* genotypes and the WT (Fig. **4b**). As a result of the vio→zea exchange, we observed a reduction in the contribution of xan to far-red emission, particularly evident in the 510-540 nm range, similar to what observed for the *a*603-NH mutant (Fig. **S19**). The xan exchange also led to a slight change in the chl a/b ratio, accounting for enhanced absorption at 465 nm and 652 nm (Fig. **S20a**), consistent with previous observations (Ballottari *et al*., 2014) (Fig. **S20d**).

To analyze pigment-pigment interaction in WT and mutant LHCI complexes, we recorded CD spectra in the visible region (350-750 nm). In the Q_Y_ region, the spectra of all genotypes displayed signature peaks typical of LHC (i.e., -/+/-), indicating a similar and conserved structural conformation and pigment organization (Mozzo *et al*., 2008). The CD spectra of the WT and *a*603-NH complexes revealed a large difference in the far-red region (λ > 700 nm), where the mutant showed a markedly reduced (-) signal associated with the excitonic interaction responsible for the RF (Fig. **4c**). The difference spectrum (black line) evidenced the disappearing of a (-) low-energy band in *a*603-NH LHCI (690-730 nm), which was compensated by a high-energy (+) band appearing at 684 nm. This conservative signal aligns with the loss of excitonic interactions between chl *a*603 and *a*609, as previously reported (Morosinotto *et al*., 2003; Wientjes *et al*., 2012).

In the Soret region, interpreting the CD spectra is complicated due to the superimposition of signals from chls and cars. However, we detected notable differences between 465-530 nm when comparing both the *a*603-NH and *npq2* samples with the WT. These differences can be attributed to genuine changes in pigment-pigment interactions, variations in Xan composition (vio *vs*. zea) or minor losses of cars during purification (Fig. **S20d**).

Our observation suggested a minor, yet significant, role of the xan at L2 in interacting with the chl a603-a609 pair, thus contributing to modulating the overall absorption of the cluster towards longer wavelengths.

### Excited state calculations

To model the effect of the *a*603-NH mutation on the red-shifted absorption at the molecular level, we performed structure-based polarizable quantum mechanics/molecular mechanics (QM/MM) calculations of excited states on the Lhca4 WT and *a*603-NH mutant models, including charge-transfer excitations (Fig. **5a**). In the spectra simulated with the standard exciton model (noCT in Fig. **5c**, Fig. **S21**), neither the WT nor the a603-NH exhibited a signature of the RF observed in the experiments, notably the broad band peaking at >700 nm. Conversely, by adding the CT contribution for the *a*603-*a*609 dimer in the calculations, we observed a low energy band (∼710 nm) in the simulated spectrum of the WT (Fig. **5b**), resembling the experimental observations in the isolated rLhca4 (Wientjes *et al*., 2012). In contrast, virtually no change in the spectrum was observed for the mutant (Fig. **5c**). This indicates that the major contribution to the WT/*a*603-NH absorption shift resides in the different extent of the coupling of CT states to local excitations between chl *a*603-*a*609, supporting a previous proposal (Wientjes *et al*., 2012; Sláma *et al*., 2023).

**Figure 5.**
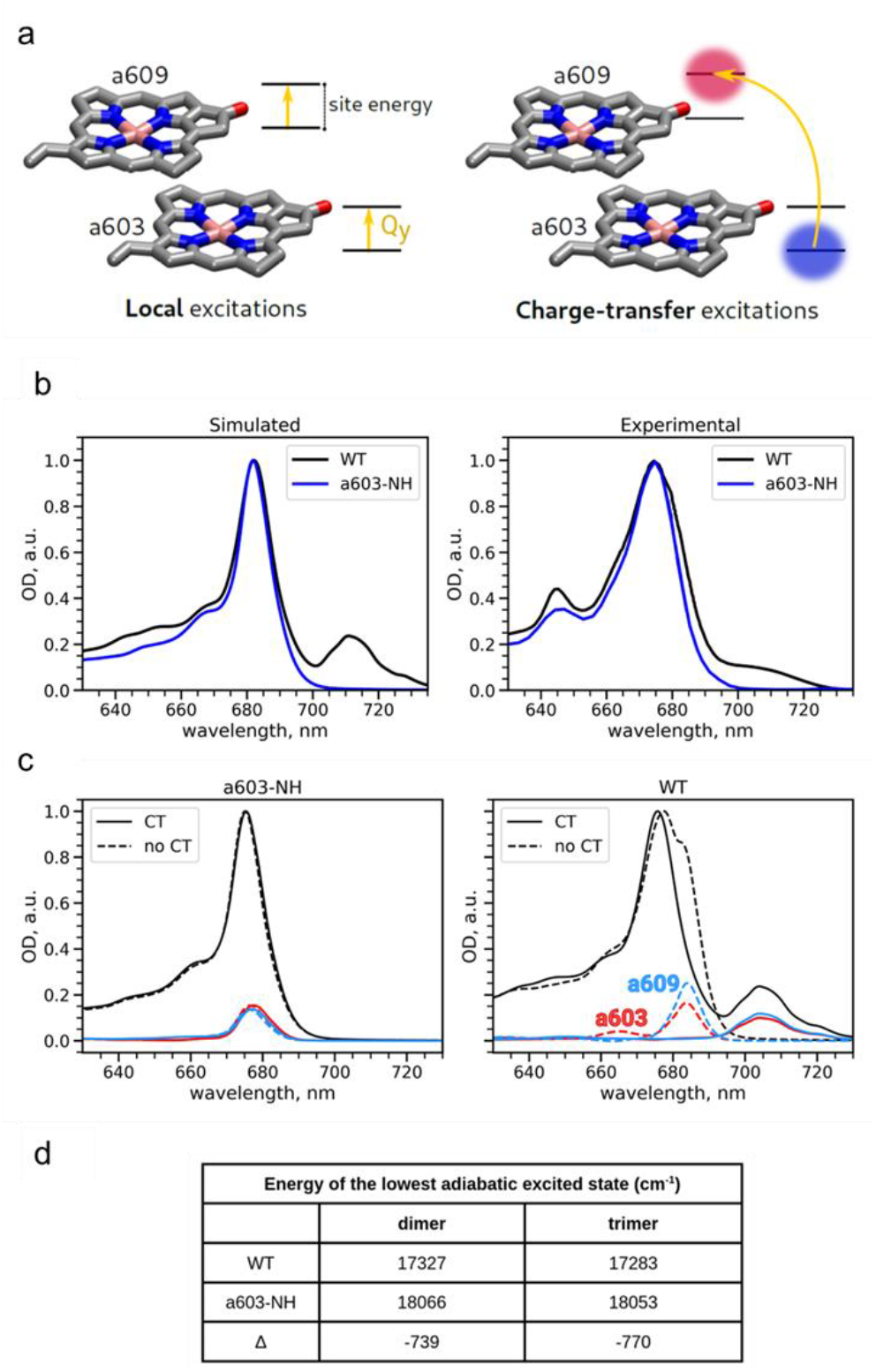
(a) Schematic depiction of local excitations (left) and charge transfer excitations (right). In local excitations, electron transitions occur within orbitals of the same molecule; in charge-transfer excitations electrons are promoted from orbitals localized on one molecule to orbitals localized on another molecule. (b) Comparison of WT and *a*603-NH absorption spectra in the simulations (left) and experiments for isolated Lhca429 (right). (c) Effect of adding CT excitations to the exciton model in the *a*603-NH mutant (left) and WT (right). (d) Energy of the lowest excited state in the *a*603-*a*609 dimer and in the *a*603-*a*609-vio trimer. The experimental spectra of WT and a603-NH are adapted with permission from Ref. (Wientjes *et al*., 2012). Copyright © 2012 Elsevier B.V. All rights reserved.

A secondary effect can be noticed in the WT spectrum simulated without CT states (Fig. **5c**), namely the appearance of a red-shifted shoulder, corresponding to *a*603-*a*609 absorption. While this shoulder is clearly more blue-shifted than the red band at ∼710 nm, it indicates red-shifted site energies for *a*603-*a*609, in contrast with the *a*603-NH mutant. Nonetheless, only with CT states does the model predict the significantly red-shifted band observed in the experiment.

We also assessed the electronic interactions between the chls dimer and L2 vio. To this end, we computed the excited states for two supermolecules, one formed by *a*603 and *a*609 (dimer), and the other formed by the same two chls and the vio (trimer). We investigated the energy of the lowest excited state responsible for fluorescence emission, to explore the effect on the electronic structure of including or excluding vio (Fig. **5d**). The small change observed (15 to 50 cm^−1^) is consistent with the varying treatment of the car (as point charges in the dimer *vs*. an active molecule in the trimer), indicating that Vio in L2 does not effectively mix its orbitals with the two chls and therefore does not significantly contribute to CT excitation within the *a*603-*a*609 pair. The change in the difference between the lowest dimer and trimer energy states in WT and a603-NH (13 and 44 cm^−1^, respectively) suggests a very minimal involvement of the car in L2 in tuning the low-energy absorption of the cluster, even smaller than what our experimental results indicate (Fig. **4**).

### The role of the extra chromophore chl *a*615

Lhca3 and Lhca4 bound an additional chl molecule, *a*615 (referred to as chl *a*617 in (Qin *et al*., 2015)), which was coordinated by a His residue located on either the third or fourth turn of the C helix of Lhca3 and Lhca4, respectively (Fig. **3**). This pigment was absent in Lhca1 and Lhca2, leading to speculation about its involvement in RF formation (Melkozernov & Blankenship, 2003) (Fig. **3**), as the positioning of chl *a*615 allowed for favorable dipolar coupling with chl *a*609. Notably, the EC value between chl *a*615 and chl *a*609 was 79 cm^−1^ for Lhca3, whereas it dropped to 19 cm^−1^ in Lhca4 due to the differing orientation of the chlorin ring in relation to chl *a*609. Although His residues were also present in the second turn of the C helix of Lhca1 and Lhca2, the lack of electronic density from cryo-EM suggested that chl *a*615 was absent in these subunits.

An additional lut molecule was located near the chlorin ring of chl *a*615 in Lhca4 (Fig. **S22a**), positioned to allow strong dipolar coupling (492 cm^−1^) with this chl. This lut had previously been observed only in the cryo-EM structure of *Z. mays* PSI-LHCI (PDB 5ZJI (Pan *et al*., 2018)) and in the X-ray structure of *P. sativum* PSI-LHCI (Qin *et al*., 2015). The xan was situated at the interface between Lhca1 and Lhca4, in contact with both subunits, and one of its hydroxyl groups formed a hydrogen bond with the carbonyl oxygen of Ser210 in Lhca1 (Fig. **S22b**). Consequently, it might got lost during the purification of monomeric Lhca4.

To investigate the potential role of chl *a*615 in far-red light absorption, we generated mutants lacking this chromophore in both Lhca3 and Lhca4 by substituting the His-binding residue with non-binding Ala (*a*615-H→A) or Ile (*a*615-H→I) (Remelli *et al*., 1999; Guardini *et al*., 2022).

When recording 77 K emission spectra from intact leaves of genotypes with and without chl *a*615, we observed a small blue-shift from 736 nm to 733 ± 1 nm (Fig. **6a, b**). However, in the isolated PSI-LHCI complex, both genotypes showed the same emission at 734 nm (Fig. **5c**). The ∼2-3 nm red-shift in leaf samples compared to isolated supercomplexes can be attributed to self-absorption effects (Weis, 1985). The *a*603-NH mutant and *koLhca3 koLhca4* exhibited a blue-shifted λ_max_ to ∼724 nm, similar to the emission of the PSI core complex (Croce *et al*., 1998). We concluded that chl *a*615 does not play a role in forming RF. To explain the 3 nm shift observed in leaf emission with and without chl *a*615, we analyzed the pigment-protein organization of thylakoids by sucrose gradient ultracentrifugation. Fig. **S23** compares the fractionation patterns from solubilized thylakoids from the WT and chl *a*615-less mutants. In addition to the lowest (higher MW) band containing the fully assembled PSI-LHCI supercomplex, a prominent green band containing the PSI-core complex was present in both *a*615 mutant lines, while it was very faint in the WT (Wientjes *et al*., 2009). Consistently, the upper band (with the lowest MW) was enriched in the mutants compared to the WT. We interpreted these results as indicating a de-stabilization of the dimeric Lhca1-Lhca4 and Lhca2-Lhca3 dimers consequent to the missing chl *a*615. This was further confirmed by a second sucrose gradient fractionation upon treating the isolated PSI-LHCI with Zwittergent-16, a procedure that allows the isolation of LHCI dimers from the PSI core (Fig. **6d**). The pattern from the WT yielded both monomers and dimers of Lhcas, while only monomers were observed in both chl *a*615 mutant lines. These results support the hypothesis that chl *a*615 plays a key role in stabilizing the dimeric Lhca complexes.

**Figure 6.**
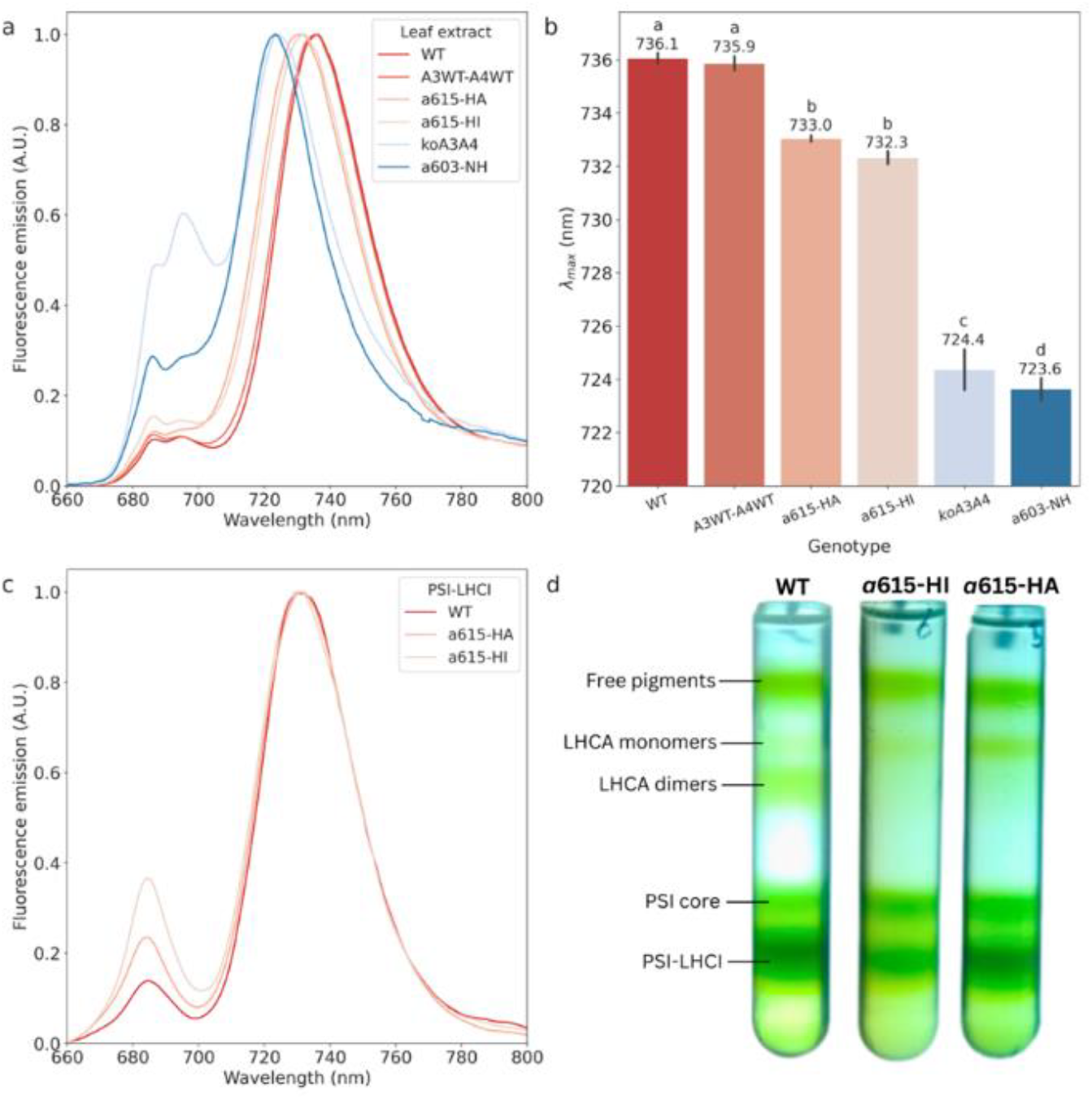
Spectral characteristic of different chl-binding mutants of A. thaliana. (a) Fluorescence emission spectra (λ_exc_ in nm) measured on leaf extracts for different genotypes of *A. thaliana*: WT, *koLhca3 koLhca4* lines complemented with WT sequences of Lhca3 and Lhca4 (A3WT-A4WT), mutants lacking chl *a*615 (a615-HA and a615-HI), *koLhca3 koLhca4* (koA3A4), and *a*603-NH mutant (*a*603-NH), and normalized to the λ_max_. (b) Barplot of the peak emission wavelength (λ_max_), measured on the same genotypes. The values on the individual bars represent the mean λ_max_ in nm, the error bars correspond to the standard deviations (n = 5 - 10). Values that are significantly different (ANOVA followed by Tukey’s post-hoc test at a significance level of P < 0.05) are marked with different letters. c) PSI-LHCI Fluorescence emission spectra (λ_exc_ in nm) measured on the WT, *a*603-HA and *a*603-HI genotypes. d) Sucrose gradient fractionation of solubilized PSI-LHCI from WT, *a*603-HA and *a*603-HI plants. Five pigment-containing bands were resolved and identified as: free-pigments, monomeric LHCA, dimeric LHCA, PSI-core complex, and PSI-LHCI supercomplex. Experiments in panels c and d were repeated independently twice, with similar results.

## DISCUSSION

A major trend in the evolution of PSI-LHCI within the green lineage has been the progressive increase in the amplitude of the far-red tail of the absorption spectra. This change is consequent to the appearance of low-energy absorption forms in LHCI, developed around 489-403 MYA, likely in response to the far-red enriched radiation made available by the development of vegetation canopies (Fig. **1**).

Notably, these variations in spectral properties occurred despite the structural similarities among their LHC subunits (Iwai *et al*., 2024). This can be observed through the structural superposition of the red clusters from representative C3 (*A. thaliana* and *F. verschaffeltii*), C4 (*Z. mays*), green algae (*C. reinhardtii*), and mosses (*P. patens*) Lhca4 and Lhca3 (Fig. **7**), or the corresponding homologous subunits. The substitution of Asn, coordinating chl *a*603, with a His residue is a structural feature known to disrupt the excitonic interactions that generate the low-energy states in *At*-rLhca3 and *At*-rLhca4 (Morosinotto *et al*., 2003; Wientjes *et al*., 2012; Li *et al*., 2023). However, the relationship between Asn ligand and the presence of Lhca-associated RFs appears overly simplistic; indeed, in higher plants which exhibit the strongest far-red emission (i.e., *A. thaliana, Z. mays*, and *F. verschaffeltii*), chl *a*603 is coordinated by Asn residues in both Lhca3 and Lhca4, while in *C. reinhardtii* chl *a*603 is coordinated by a His residue in Lhca3 and Lhca8 (the Lhca4 homologue). Notably, the Asn ligand is present in other loosely bound Lhca subunits (Stauber *et al*., 2009; Su *et al*., 2019) which are located in more distal positions relative to the PSI core complex (Huang *et al*., 2021). Moreover, *P. patens*, which occupies an intermediate position both evolutionary and in terms of absorption properties, has an Asn residue that coordinates chl a603 in Lhca3, but not in Lhca2b, its Lhca4 structural homologue.

**Figure 7.**
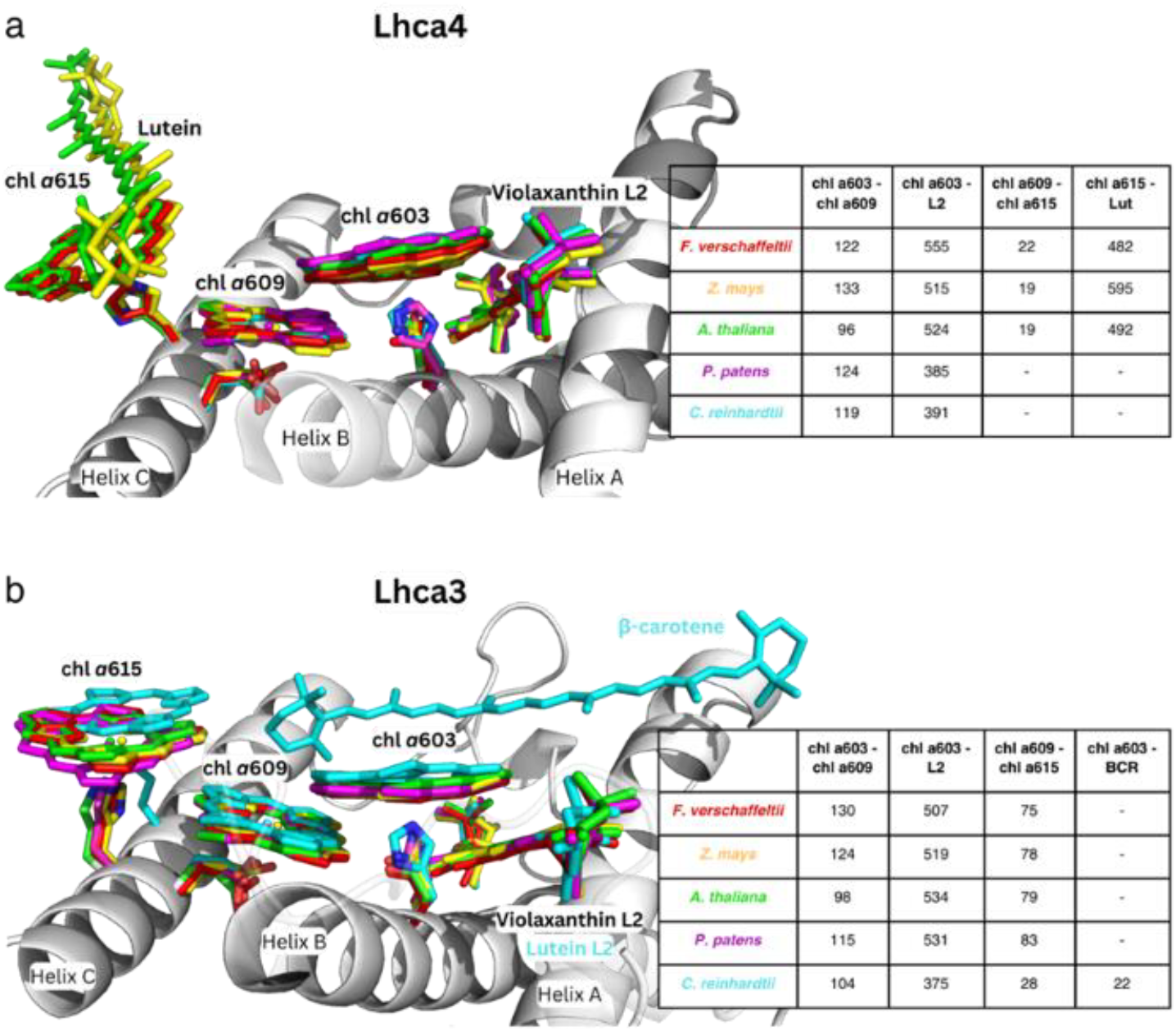
Structural superposition and excitonic coupling analysis (values in cm^−1^) of the red cluster pigments from *F. verschaffeltii* (PDB 8WGH), *Z. mays* (PDB 5ZJI), *A. thaliana, P. patens* (PDB 7XQP), and *C. reinhardtii* (PDB 7ZQC). The analysis focused on (*a)* Lhca4 or the corresponding subunit, i.e. Lhca8 for *C. reinhardtii* and Lhca2b for *P. patens* (Yan *et al*., 2021; Gorski *et al*., 2022) and (*b)* Lhca3.

Interestingly, Asn remains the coordinating residue for chl *a*603 in the seagrasses *P. oceanica* and *C. nodosa*, despite their loss of RF after returning to the marine environment (Fig. **1**). This observation reinforces the idea that the nature of the chl *a*603 ligand is not the sole factor determining the presence of RF.

The PSI-LHCI *a*603-NH supercomplex showed a significant absorption reduction in the >700 nm region, along with a ∼13 nm blue-shifted fluorescence emission at cryogenic temperatures (Fig. **2**). This shift is notably larger than that observed in genotypes with the *a*603-NH mutation limited to the Lhca4 antenna (Li *et al*., 2023). In the WT PSI-LHCI (Wientjes *et al*., 2009), the terminal emitter is the antenna complex, which emits at at 734 nm. In contrast, the *a*603-NH mutant has the PSI core complex as the emitter, at 721 nm (Ihalainen *et al*., 2003; Akhtar & Lambrev, 2020). This difference complicates the assessment of the real extent of the emission blue-shift caused by the mutation in LHCI. This was possible upon purification of LHCI from WT and *a*603-NH mutant (Fig. **2**), allowing for the measurement of a 39 nm blue-shift between the peak emissions. This finding aligns with previous studies on *in vitro* reconstituted complexes, which reported a shift of ∼45 nm (Morosinotto *et al*., 2003; Wientjes *et al*., 2012).

By comparing the cryo-EM structures of the WT and *a*603-NH mutant PSI-LHCI, we detected a ∼0.7 Å shift in the chlorin ring of chl *a*603, induced by the Asn→His substitution in Lhca3 and Lhca4 structures (Fig. **3**, **S13**). Although this structural difference is subtle, its impact on the coupling with charge-transfer (CT) states that underlie RF formation could be considerable, given the sensitivity of these interactions to distance (Cupellini *et al*., 2018). To connect the effect of this structural shift with the observed spectroscopic properties, we performed EC calculations using the point-dipole approximation (Liguori *et al*., 2015). While the Asn→His substitution changed the separation between the chlorin rings of chl *a*603 and chl *a*609, it also increased the distance between the chl *a*603 chlorin ring and the neighboring vio L2. This change led to a substantial change in the chl *a*603–vio L2 coupling value, which dropped from ∼530 cm^−1^ in the WT to ∼380 cm^−1^ in the *a*603-NH mutant for both Lhca3 and Lhca4 subunits (Fig. **3**).

We noticed that the chl *a*603–vio L2 coupling value serves as a good indicator of the presence of RF (Fig. **3**, **6**, **S22**). Species lacking RF (*C. reinhardtii*) showed values <400 cm^−1^, while species with the highest red absorption properties (*F. verschaffeltii, Z. mays*, and *A. thaliana*) have values >500 cm^−1^. Interestingly, *P. patens*, which has red absorption properties that are intermediate between those of plants and algae, has EC values >500 cm^−1^ only in Lhca3. Moreover, the strength of the chl *a*603–vio L2 coupling correlates well with the fluorescence emission λ_max_ (Fig. **1**).

Based on this structural and computational analysis, we investigated the influence of the xan in position L2 on low-energy absorption, by applying two investigation strategies. First, we compared 77 K excitation spectra of the *a*603-NH and WT PSI-LHCI (Fig. **4**) and observed a systematically reduced intensity (by 6%) in the 455-505 nm region of the *a*603-NH PSI-LHCI supercomplex, an effect that can be attributed to the contribution of cars (Ashenafi *et al*., 2023). Second, we analyzed the PSI-LHCI supercomplex from the *npq2* genotype, in which the L2 vio binding site in PSI-LHCI supercomplex is occupied by zea (Ballottari *et al*., 2014). The structural difference between the two xan caused a ∼2 nm blue shift in *npq2* compared to WT 77 K fluorescence emission spectra and a concomitant reduction in the amplitude of the red-emission tail at 752 nm (Fig. **S20**). Interestingly, in the 480-510 nm region, where xan absorption is prominent, the PSI-LHCI from *npq2* showed a lower contribution to far-red fluorescence emission compared to WT PSI-LHCI (Fig. **4b**), consistent with the amplitude of the 516 nm CD signal (Fig. **4d**). Using QM/MM analysis of the excited states of the dimer *a*603–*a*609 and the trimer *a*603–*a*609–vio(L2) on the WT and *a*603-NH experimental structures of Lhca4 we show that the local rearrangement of the pigments *a*609– *a*603 due to the *a*603-NH substitution impacts the electronic overlap between orbitals (Fig. **S25**), leading to a decrease in CT couplings that are responsible for the low-lying red-states in the WT (Fig. **5b**).

Excited state calculations showed that the standard excitonic model was insufficient to explain the red-shifted absorption of the pigment cluster, while the introduction of CT states was needed to best simulate the experimental absorption spectra (Fig. **5c**).

While the inclusion of vio(L2) in the computed CT states has only a minor contribution to the far-red absorption (Fig. **5c**).

Our structural, spectroscopic and multiscale excited-state calculations analysis, suggests that while the origin of the low-energy absorption in PSI-LHCI and of RF lies in the CT states of the chl a603–a609 pair, the xan in L2 may play a minor role in modulating the energy transfer towards the chl a603–a609 pair, thus contributing to the extension of the absorption of far-red photons.

One additional chl, chl *a*615 was located close to the “red cluster” specific of the Lhca3 and Lhca4 isoforms of higher plants (Melkozernov & Blankenship, 2003). Chl *a*615 exhibited EC values with chl *a*609 similar to the canonical red pair chl *a*603–*a*609, especially in the Lhca3 isoforms due to the more favorable orientation (Fig. **7**). Upon site-directed mutagenesis to selectively remove chl *a*615 in both Lhca3 and Lhca4, low-temperature fluorescence emission spectra allowed ruling out the involvement of Chl *a*615 in RF. Nevertheless, this experiment showed that chl *a*615 in Lhca4 was needed for stabilization of the Lhca1-Lhca4 dimer and/or its assembly with the PSI-core (Fig. **6d**), (Fig. **S23**). Taken together, our detailed analysis of the large 3-chl-1-xan pigment assembly in the LHCI domain expressing RF highlights the role of the charge-transfer states within the *a*603–*a*609 pair (Sláma *et al*., 2023). Xan L2 effect was determined to be of minor importance despite undergoing the largest structural change when comparing WT vs a603-NH Lhca3 and Lhca4. Finally, we report on two chl a615 and one lut ligand, which, although not contributing to the formation of RF, were proven to substantially contribute to the assembly of LHCI dimerization and assembly with PSI-core complex.

Further research will be necessary to better clarify the contribution of the protein environment surrounding the “red cluster,” for example, by comparing high-resolution structures of red-shifted and blue-shifted Lhcas such as from *F. verschaffeltii* and seagrasses.

Tuning chl absorption towards spectral regions where photons are available, especially under dense canopies, is a privileged strategy for enhancing crop photosynthetic yield. The detailed knowledge of structure-function relations within LHC antenna proteins will contribute to the rational engineering of crops (Ort *et al*., 2015; Cutolo *et al*., 2023).

## ACKNOWLEDGMENT

RB acknowledges financial support from the European Research Council (ERC Advanced Grant 101053983-GrInSun). Part of this work was carried out at Unitech NOLIMITS, an advanced imaging facility established by the University of Milan. The authors acknowledge Dr. Gabriele Procaccini from Anton Dohrn Experimental Marine Station (Naples) for providing *P. oceanica* and *C. nodosa* plants.

## AUTHOR CONTRIBUTIONS

R.B., S.C. and L.D. conceived the work and designed the experiments. Z.G., V.F.P. and A.A. carried out the construction of mutants and performed their biochemical and spectroscopical characterization. Z.G., S.C. and A.A. carried out the preparation of the samples for cryo-EM. D.M.V.B. e A.C-S. conducted cryo-EM sample preparation and data collection. S.C. and D.M. analyzed cryo-EM data and reconstructed PSI-LHCI structures. D.M. performed bioinformatics analysis and EC calculations. E.B., C.J., L.P-G., L.C. and B.M. carried out quantum chemical calculations and analyses. D.M., S.C., and Z.G. wrote the original text draft, and all authors discussed the results and contributed to drafting the manuscript.

## DATA AVAILABILITY

Sequence data from this article can be found in the Arabidopsis Genome Initiative under accession numbers At1g61520 (LHCA3), At3g47470 (LHCA4) and At5g67030 (ZE). The KO lines were obtained in the NASC under stock numbers N876497 (*koLhca3*), N679009 (*koLhca4*) and N60000 (*npq2*).

The Cryo-EM maps and coordinates have been deposited in the EMDB and wwPDB, respectively: PSI–LHCI WT (cryo-EM map, EMD-51219; consensus refinement map; PDB: 9GBI), PSI–LHCI *a603-NH* mutant (cryo-EM map, EMD-51227; PDB: 9GC2).

## SUPPORTING INFORMATION

Additional Supporting Information may be found online in the Supporting Information section at the end of the article. **Supplemental methods. Figure S1:** Sample preparation and characterization. **Figure S2:** Cryo-EM data processing workflow for AtPSI-LHCI WT. **Figure S3:** Cryo-EM data processing workflow for AtPSI-LHCI *a603-NH*. **Figure S4:** Spectral characteristic of different of *A. thaliana* WT and *a603-NH* mutant. **Figure S5:** Absorption and emission spectra of PSI core complex of *A. thaliana* WT and a603-NH mutant. **Figure S6:** Atomic models of PSI-LHCI subunits and selected ligands superimposed on cryo-EM maps. **Figure S7:** Top and side view of the PSI-LHCI WT supercomplex adn pigment organization. **Figure S8:** Positions of ligands in the PSI-LHCI WT of *A*.*thaliana*. **Figure S9:** Lipid molecules found in PSI-LHCI supercomplexes. **Figures S10:** Superposition of the PSI WT from *A. thaliana* (PDB 9GBI, white) and from P. sativum (PDB 7DKZ) **Figure S11:** Global superposition RMSD of the AtPSI-WT (PDB 9GBI) structure with PSI-WT structures from PDB 8J7B, 8JZA (both from cryo-em data), and 7DKZ. **Figure S12:** Position of chl, carotenoids and lipid molecules in PSI-LHCI supercomplexes. **Figure S13:** Pigment content of the LHCI antenna subunits of *A. thaliana* WT (PDB 9GBI). **Figure S14:** Atomic models of the “red cluster” pigments in Lhca3 and Lhca4. **Figure S15:** Pigment distances in the “red cluster” of Lhca3 and Lhca4. **Figure S16:** Top and side view of superimposed structures of chl a603, chl a609, chl a615, Violaxanthin L2 and Lutein. **Figure S17:** Top and side view of superimposed structures of chl a603, chl a609, chl a615, Violaxanthin L2 and Lutein. **Figure S18:** Excitonic coupling absolute values between pigments in LHCI WT and a603-NH. **Figure S19:** Difference spectrum between RT absorption and 77 K excitation spectra in the 425-690 nm region of PSI-LHCI from *A. thaliana* WT and a603-NH. **Figure S20:** Absorption and emission spectra of PSI-LHCI complex of A. thaliana WT and and *npq2* mutant, absorption spectra and pigment composition of LHCI complexes of *A. thaliana* WT and and *npq2* mutant. **Figure S21:** Site energies of Lhca4 chls for WT and a603-NH mutant. Absorption spectra of WT and mutant simulated using the simple exciton model. **Figure S22:** Structural superposition of Lhca4 WT from *A. thaliana* (PDB 9GBI,), *P. sativum* (PDB 5L8R), *Z. mays* (PDB 5ZJI). Cryo-em map for Lut over chl a615 in Lhca4. **Figure S23:** Sucrose gradient fractionation of thylakoid membranes of WT and *a*615 mutant lines. **Figure S24:** Superposition of Lhca4, Lhca2 from *A. thaliana* WT (PDB 9GBI) and Lhcb4 from *P. sativum* (PDB 5XNL) and excitonic couplings between pigments pairs in the red cluster. **Figure S25:** Visual representation of the overlap between LUMO orbitals of chls *a*603 and *a*609 as computed for the WT and *a*603-NH structures. **Figure S26:** Sequence alignment of the sequences of the synthetic genes encoding for Lhca3/a4_WT and Lhca3/a4_a615-HA and Lhca3/a4_a615-HI. **Table S1:** AtPSI-603-NH structural model. CLA, chl *a*; CHL, chl *b*; BCR, β-carotene; LUT, lutein; XAT, violaxanthin; DGD, digalactosyl-diacyl glycerol (DGDG); LMG, 1,2-distearoyl-monogalactosyl-digliceride; LHG, 11,2-dipalmitoyl-phosphatidyl-glycerol; LMT, dodecyl-β-D-maltoside; PQN, phylloquinone; SF4, Fe4-S4 cluster. **Table S2:** List of the primers used to obtain and characterize the *a603-NH* mutant lines. **Table S3:** Cryo-EM data collection, refinement, and validation statistics. **Supplemental References**

