## supplemental methods figures tables for "Structural determinants for red-shifted absorption in higher-plants Photosystem I"

ADDITIONAL METHODS:

**Assembly of complementation constructs**

To generate the *a603-NH* mutant, *LHCA3* and *LHCA4* genes were PCR-amplified from *Arabidopsis* WT (col-0) genomic DNA. The amplified fragments, which included the 5’-UTR and 3’-UTR regulatory regions along with the coding region of *LHCA3* or *LHCA4*, were obtained using the following primer pairs:

Lhca3_FW 5’-GGGGACAAGTTTGTACAAAAAAGCAGGCTTCTGAAGCAGATACTAATCTAATCTGAGC-3’;

Lhca3_RV 5’-GGGGACCACTTTGTACAAGAAAGCTGGGTCAAGGTGGTTTGGGAGATACAGA-3’

Lhca4_FW 5’-GGGGACAAGTTTGTACAAAAAAGCAGGCTTCTTTTGCATGGTGATTTCCAG-3’

Lhca4_RV 5’-GGGGACCACTTTGTACAAGAAAGCTGGGTCTCCTCAATCAGGTTGGCTTA-3’

These primers were designed to include attB clamps for the Gateway cloning system (Invitrogen). The resulting amplicons were cloned into pDONR221 vectors and subsequently recombined into pK7WG2 and pH7WG2 plant destination vectors (Karimi et al., 2002) for *LHCA3* and *LHCA4*, respectively. Specific primers (Table **S2**) and the QuickChange Stratagene kit were used to substitute Asn encoding triplets with His triplets. The resulting pK7WG2-*LHCA3*-N103H and pH7WG2-*LHCA4*-N99H vectors were co-transformed into *koLhca3 koLhca4* background, and transformant lines were selected on MS-agar medium supplemented with 50 μg/mL kanamycin and 25 μg/mL hygromycin.

To produce *koLhca3 koLhca4* complemented with either WT *(A3WT-A4WT*) or *a615-HA* and *a615-HI* mutations, the 5’-UTR, coding sequence and 3’-UTR regions were sequence-optimized for the Golden Braid modular cloning system (Vazquez-Vilar *et al.*, 2017) and obtained as separate synthetic sequences from Genewitz (www.genewiz.com) (Fig. **S26**). The final transcription units encoding for each gene were assembled using type IIS restriction enzymes BsaI and BsmbI (Sarrion-Perdigones *et al.*, 2013). The resulting pDGB3α1-*LHCA3*_H168A-*LHCA4*_H151A and pDGB3α1-*LHCA3*_H168I-*LHCA4*_H151I plant expression vectors were used to complement *koLhca3 koLhca4* plants, and transformant lines were selected on MS-agar medium supplemented with 50 μg/mL kanamycin.

**Gel electrophoresis**

SDS-PAGE analysis was performed using the Tris-Glycine buffer system (Laemmli, 1970) with the modifications described in (Ballottari *et al.*, 2004).

**Phylogenetic analysis**

The Timetree in Figure 1a was obtained using TimeTree5 (Kumar *et al.*, 2022).

Fluorescence emission spectra and the fluorescence emission maxima (Fig. **1b**) were recorded at cryogenic temperature. The experimental procedure was as follows: leaf tissue was frozen in liquid nitrogen, ground to a fine powder, and resuspended in measuring buffer (10 mM HEPES pH 7.5, 20% w/v glycerol). Samples were harvested at Padua’s Botanical Garden.

*Physcomitrium patens* was grown as described by (Pinnola *et al.*, 2018). Fluorescence measurements were recorded on functional chloroplasts obtained as described in (Casazza *et al.*, 2001) upon dilution into measuring buffer.

The green algae *Chlamydomonas reinhardtii* and *Chlorella vulgaris* were cultivated in minimal media, HSM (Sueoka, 1960) and BG-11 (Allen & Stanier, 1968), respectively, until the early exponential phase. They were then diluted into measuring buffer.

For all samples, 500 μL of the suspension was directly used to record the emission spectra.

Protein sequences of *C. reinhardtii* (Cr), *P. patens* (Pp), and *A. thaliana* (At), and *F. verschaffeltii* (*F. albivenis*) Lhca3 and Lhca4 shown in Fig. **1c** were retrieved from the 7ZQC, 7XQP, 9GBI, and 8WGH PDB. For the Lhca3 and Lhca4 protein sequences of *Z. mays* (Zm) and *A. comosus* (Ac) were retrieved from Phytozome (PAC:40246382 and PAC:40271081, PAC:33052185 and PAC:33036699, respectively). *P. oceanica* and *C. nodosa* Lhca3 and Lhca4 protein sequences were obtained by a *A. thaliana* Lhca3 or Lhca4 tblastn (Altschul *et al.*, 1990) query against the organisms *Posidonia oceanica* (L.) Delile (taxid:55489) and *Cymodocea nodosa* (Ucria) Asch. (taxid:55448) (Database whole genome shotgun sequence). Multiple alignments of the protein sequences were performed with Clustal Omega (Madeira *et al.*, 2022).

Spectral deconvolution of the fluorescence traces in Fig. **1b** was performed by fitting four Gaussian curves using an in-house Python script. The initial guesses for the four central peaks were 680 nm, 690 nm, 732 nm, and 750 nm. The fraction of the area under the curve at wavelengths ≥720 nm was obtained by dividing the sum of the areas of the Gaussians above 720 nm by the sum of the areas of the four fitted Gaussians.

#### **Structure refinement for QM/MM calculations**

All calculations for the WT and *a*603-NH mutant species were performed on the cryo-EM structures presented in this work. Water molecules coordinating the chlorophyll pigments (chls) were added to the structures by homology with a previously published structure (see below). Indeed, while waters were not resolved in the density map at this resolution, their presence deeply affects the excitation energies of nearby chls and therefore must be considered. Six water molecules were added: five of them were taken from the WT structure of *Pisum Sativum* (Wang *et al.*, 2021) coordinating chls 304, 306, 307, 308 (two H-bonded water molecules); one more water molecule to coordinate the Mg center of chl 311, which was lacking a coordination residue.

Some of the mobile phytyl tails of the chls, which were not resolved in the structure, were rebuilt with the *tleap* tool of the Amber (Case *et al.*, 2020) suite and clashes with the rest of the structure were minimized by iteratively rotating the dihedral angles of the added segments through a Monte Carlo scheme. All protein residues were kept in their standard protonation, except for Mg-coordinating HIS, which were forced to be delta-protonated to ensure the metal binding. Moreover, residue Glu 146 was protonated to allow the formation of an H-bond with the carbonyl group of Chl 307, as in (Sláma *et al.*, 2023). The added segments, water molecules, and H atoms were then relaxed by MM minimization with constraints on the rest of the structure. The Amber ff14SB force-field was employed for the protein, and the TIP3P model to describe water molecules. The parameters for the cofactors Chls, 1,2-distearoyl-monogalactosyl-diglyceride (LMG), beta-carotene (BCR), lutein (LUT), and violaxantin (XAT) were taken from previous work in the group (Prandi *et al.*, 2016).

A structure refinement protocol was then applied to the prepared system to achieve a proper description of the pigment geometries and of the protein pockets. The refinement protocol comprises three steps: (1) restrained relaxation of all heavy atoms; (2) multiple cycles of simulated annealing runs; (3) QM/MM optimizations (Chung *et al.*, 2015) of the chls.

During Step 1, all the structure was relaxed with positional restraints of 10 kcal/mol-A^2^ and subsequently, in a second step, of 4 kcal/mol-A^2^ on the protein backbone atoms and on heavy atoms of XAT, LMG, LUT (only the two extrema of the conjugated chain), and finally on the internal ring connecting N atoms and on Mg for chls. Protein residues coordinating the chls were left free to move to adjust the coordination geometry. To compensate for force-field inaccuracies in describing the Mg-binding residue distances, fictitious bonds between the Mg atoms and their binding residues were added, whose equilibrium distances were assigned based on average binding distances obtained on QM/MM-optimized structures.

Step 2 consisted of 5 cycles of simulated annealing: a T ramp of 0 K - 300 K - 0 K and the Berendsen thermostat were employed. The final structure of each run was used as a starting point for the following one; the structure of the last run was finally MM-minimized. Positional restraints of 10 kcal/mol-A^2^ analogous to the ones of the previous step were used during simulated annealing to keep the overall structure close to the crystal one, while allowing the side chains to rearrange towards a more favorable conformation. In the case of the mutant structure, smaller restraints were imposed to the a603 ring to allow it to rearrange to the more hindered pocket.

Within Step 3, the chls’ internal geometry was optimized at QM/MM level using an ONIOM scheme. Each chl molecule was independently optimized; the high-level layer of the ONIOM scheme comprised the chl ring and the binding residue (protein sidechains or water molecules) and was allowed to move. The low-level layer included the protein environment within 25 A from the chl molecule and was frozen during optimization. The optimizations were run in Gaussian 16 at B3LYP/6-31G(d) level of theory (Frisch *et al.*, 2016).

The *a*603-*a*609 chl dimer was further refined together in order to achieve an accurate description of the intermolecular conformation. Indeed, it is known that CT excitations are extremely sensitive to the relative position and orientation of the pigments. A new ONIOM optimization was run, including the two chl rings and three protein residues of the pocket (His 99, Glu 154 and Arg 157) as the moving high-level layer, and the environment within 25 A as the low-level layer (kept frozen as before).

The fit of the final refined structure in the experimental density map was confirmed.

#### **Excited-state calculations**

We computed all elements (site energies and couplings) of the exciton Hamiltonian of Lhca4 on the structure refined as detailed above. The site energies of the individual chlorophylls were computed at the TD-DFT M062X/6-31G(d) level of theory. This choice is based on previous works on LHCs (Sláma *et al.*, 2020; Saraceno *et al.*, 2023). The environment effects were included through a polarizable QM/MM methodology (QM/MMPol): the QM part is composed of the chlorophyll ring (a H atom is added to saturate the valence after cutting the phytyl tail), while the rest of the protein is treated at the polarizable MM level. The charges and polarizabilities for the MM part were the same as in (Sláma et al., 2023). The chl-chl exciton couplings were calculated using the TrEsp (transition charges from electrostatic potentials) approach (Madjet *et al.*, 2006), wherein the transition charges required to compute the couplings are obtained from a fit of the electrostatic potential generated by the transition density. All QM/MMPol calculations were run using a locally modified version of Gaussian 16.

Next, to include the charge-transfer (CT) states, we built an extended Hamiltonian with CT energies and couplings. The CT states were computed for two supramolecules: (i) the *a*603-*a*609 dimer and (ii) the trimer, *a*603-*a*609-vio. For both cases, the excited states were computed on the refined structure using the Tamm−Dancoff approximation (TDA) formulation of TD-DFT at ωB97XD/6-31G(d) level of theory within the QM/MMPol scheme. ωB97XD was chosen because a good description of CT states requires the use of long-range corrected functionals. For case (i), the QM part consisted of the chlorophyll dimer, *a*603 and *a*609, while for case (ii), the QM part included vio along with *a*603-*a*609 dimer. To obtain the CT energies and the corresponding locally excited (LE) state-CT couplings from the excited state calculations, the multistate-FED-FCD diabatization scheme (Nottoli *et al.*, 2018) was employed on the first 14 excited states of the dimer. The multi-FED-FCD scheme combines the Fragment Charge Difference (FCD) and Fragment Excitation Difference (FED) methods to separate LE/CT subspaces for multiple states and extract electronic couplings. The two lowest CT states, *a*603^+^*a*609^-^ and *a*603^-^*a*609^+^, obtained from this diabatization scheme were added to the exciton Hamiltonian to build the extended Hamiltonian. The site energies and couplings in the exciton Hamiltonian corresponding to *a*603 and *a*609 are replaced by the LE energies and couplings from diabatization. To correct for the difference in level of theory between the site and CT calculations while building the extended Hamiltonian, we calculate the shift in the average of the LE state energies (*a*603* and *a*609*) from the diabatization calculation with respect to the average of the site energies and add this shift to the CT and LE energies in the extended Hamiltonian.

#### **Simulation of absorption spectra**

The spectrum of the excitonic aggregate was simulated in the disordered exciton model by summing contributions of absorption to individual exciton states. Static disorder was included as Gaussian random shifts of the pigments’ site energies and CT energies (600 realizations, with a standard deviation of 100 cm^-1^ for LE states and 1000 cm^-1^ for CT states). The homogeneous absorption lineshape of each exciton is obtained using the spectral density formalism and the second-order cumulant expansion in the so-called complex Redfield approximation (Gelzinis *et al.*, 2015). An experimentally-derived spectral density was used to describe coupling to vibrations localized on the pigment and on the protein environment (Novoderezhkin *et al.*, 2004).

All spectra were simulated using the in-house pyQME package (https://github.com/Molecolab-Pisa/pyQME).

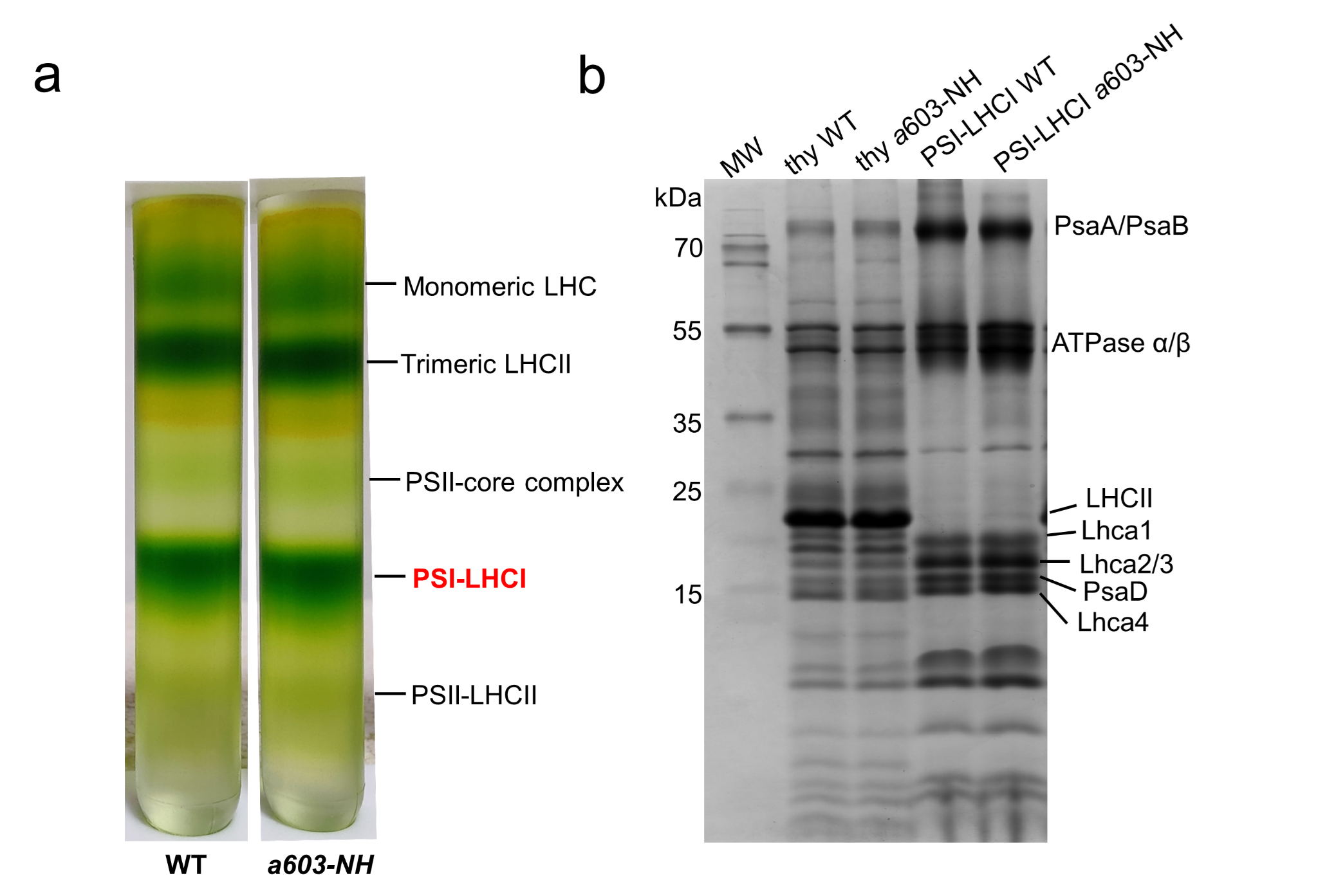

**Figure S1:** **Sample preparation and characterization.** **a)** Sucrose density gradient fractionation of solubilized thylakoid membranes from *Arabidopsis thaliana* WT and *a*603-NH mutant strains. The PSI-LHCI bands harvested for cryo-EM and further characterization are labeled in red. **b)** SDS-PAGE analysis of thylakoid membranes (thy) and PSI-LHCI supercomplexes of WT and *a*603-NH mutant.

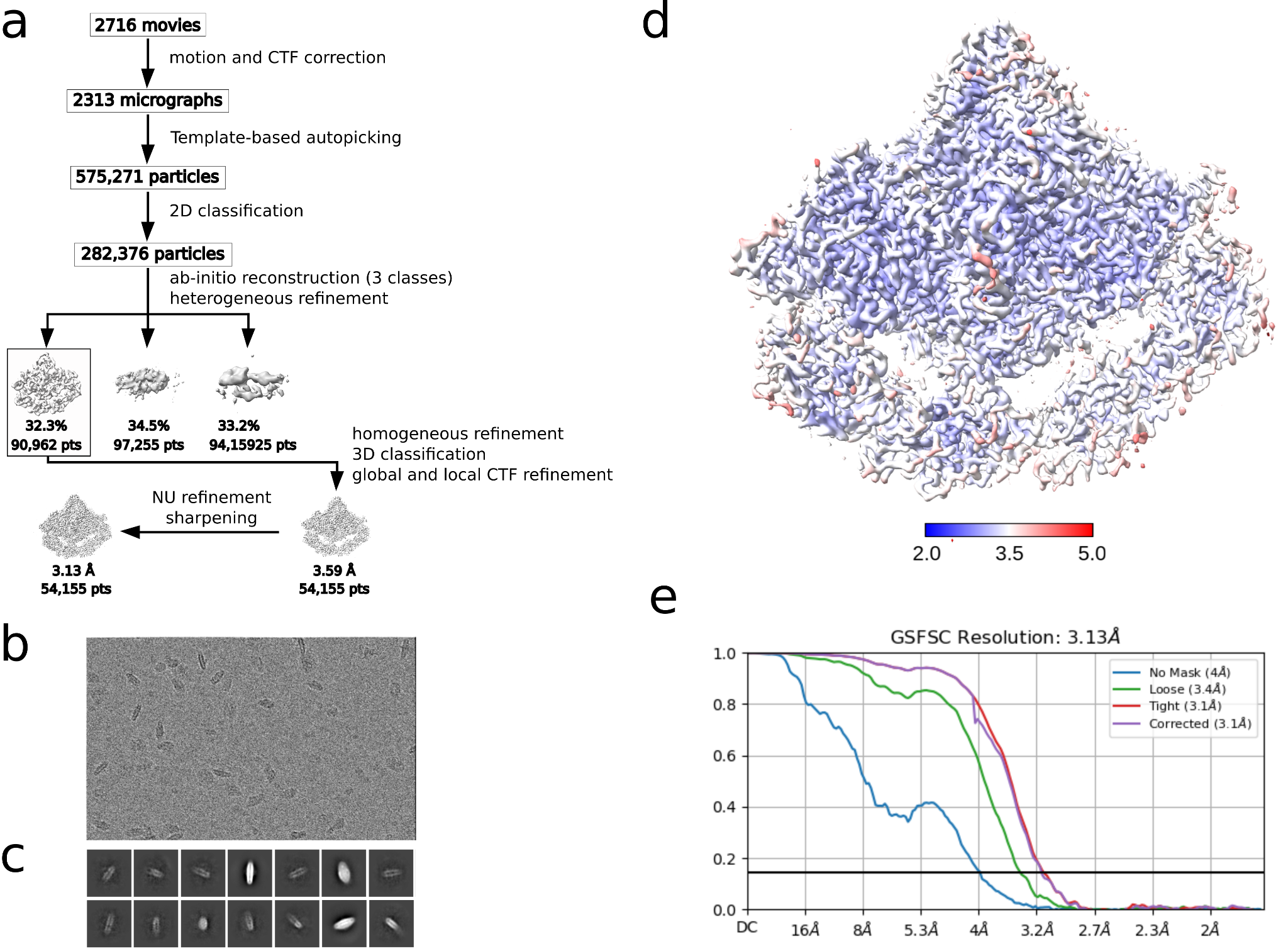

**Figure S2:** a) Cryo-EM data processing workflow for AtPSI-LHCI WT. Representative cryo-EM micrograph (b) and 2D class averages (c) of the supercomplex. d) Estimation of the local resolution of the final map. e) Gold standard Fourier shell correlation (GSFSC) curve of the final reconstruction.

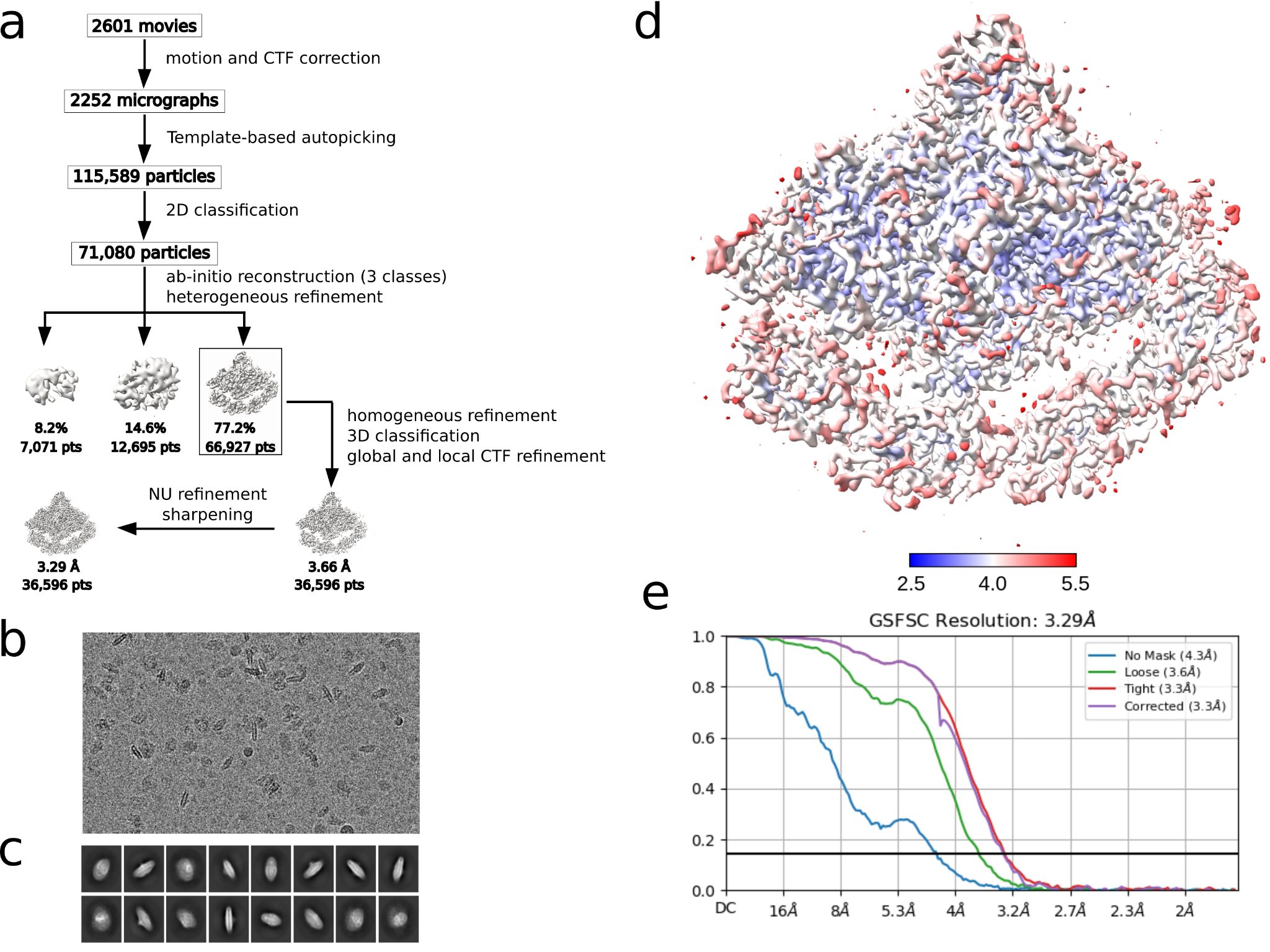

**Figure S3:** a) Cryo-EM data processing workflow for AtPSI-LHCI *a*603-NH. Representative cryo-EM micrograph (b) and 2D class averages (c) of the supercomplex. d) Estimation of the local resolution of the final map. e) Gold standard Fourier shell correlation (GSFSC) curve of the final reconstruction.Cryo-EM workflow for AtPSI-LHCI *a*603-NH.

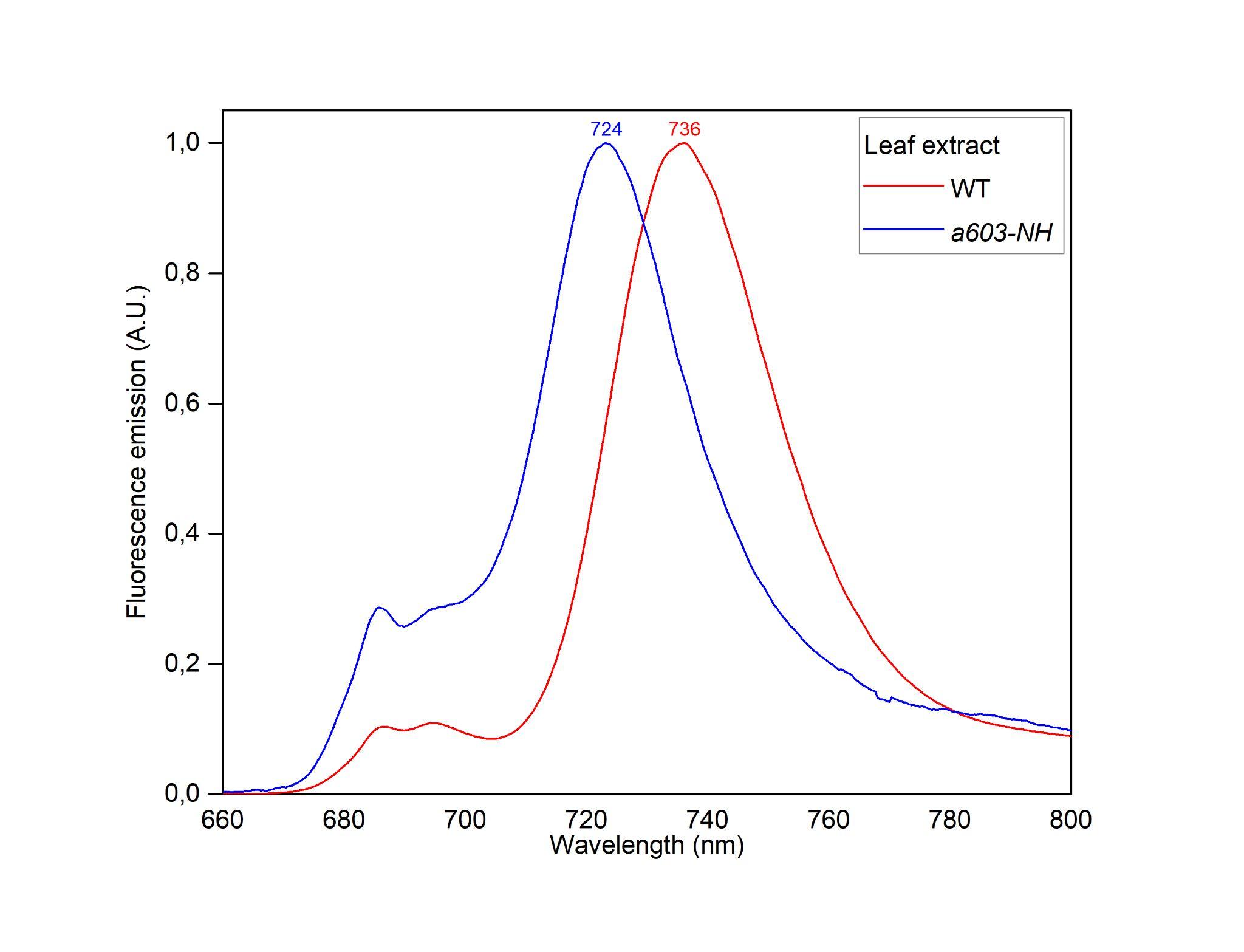

**Figure S4:** **Spectral characteristic of different of *A. thaliana* WT and *a*603-NH mutant.** Fluorescence emission spectra were measured on leaf extracts from WT, and *a*603-NH mutant (a603-NH) and normalized to the λ_max_.

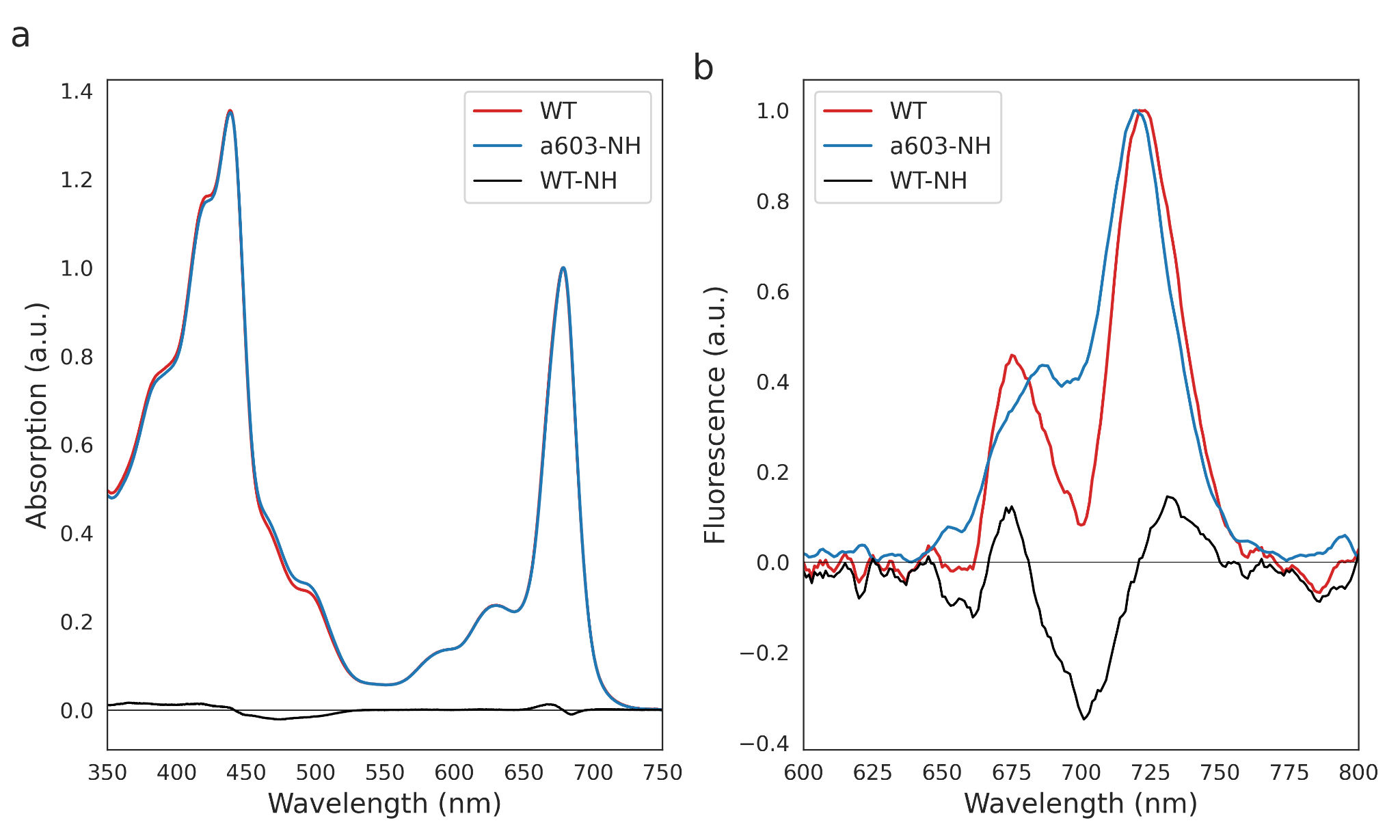

**Figure S5:** a) Room temperature (RT) absorption of the PSI core from *A. thaliana* WT and *a*603-NH. b) 77 K fluorescence emission spectra of the PSI core from *A. thaliana* WT and *a*603-NH. The WT-NH difference spectra are shown as black lines in the corresponding plots.

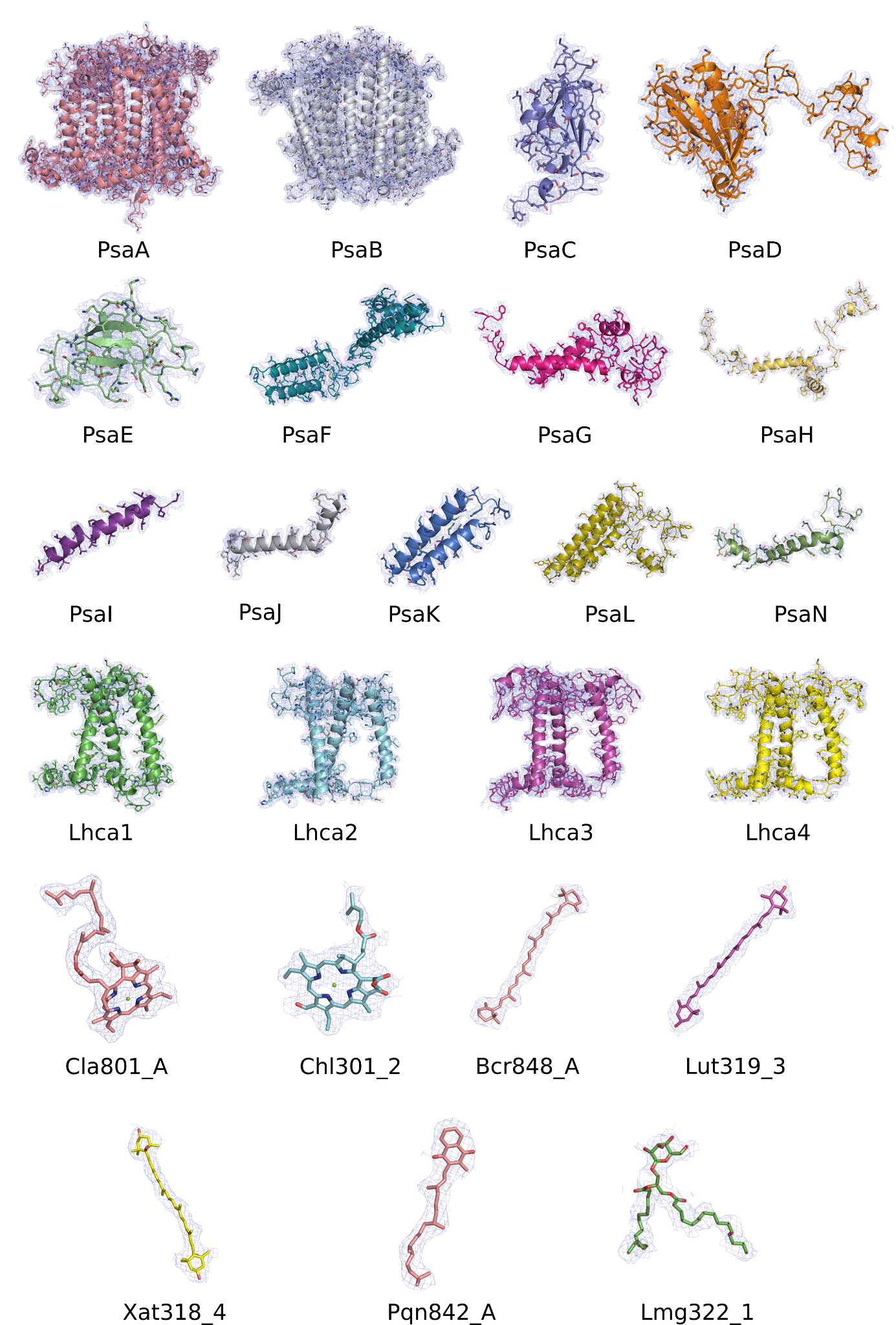

**Figure S6:** Atomic models of PSI-LHCI subunits and selected ligands superimposed on cryo-EM maps.

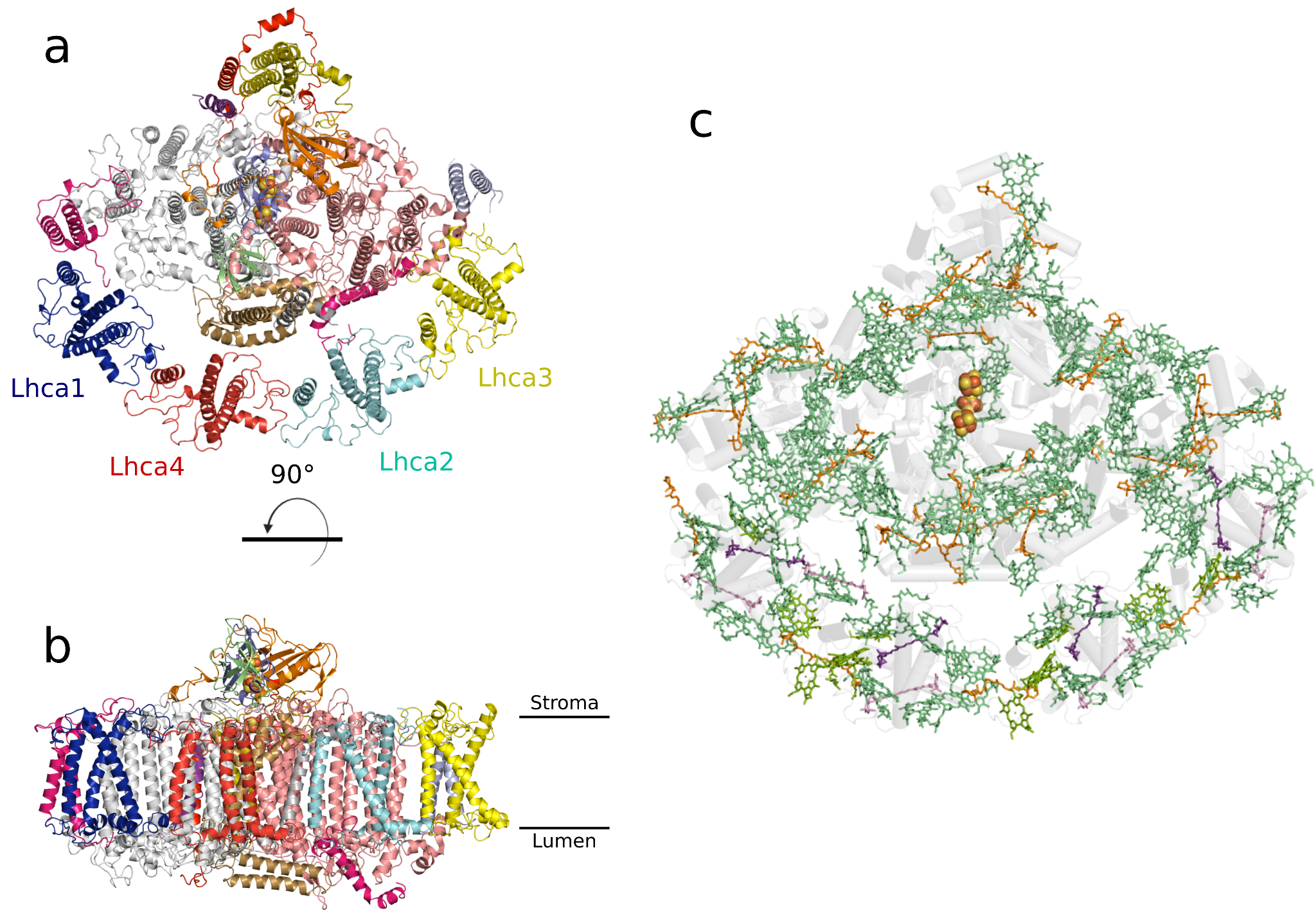

**Figure S7:** a-b) Top and side view of the PSI-LHCI WT supercomplex. Different subunits are colored with different colors. c) Pigment organization in the PSI-LHCI WT of *A.thaliana* (PDB 9GBI): chl *a* (green), chl *b* (light green), Violaxanthin (purple), β-carotenes (orange), and Lutein (pink).

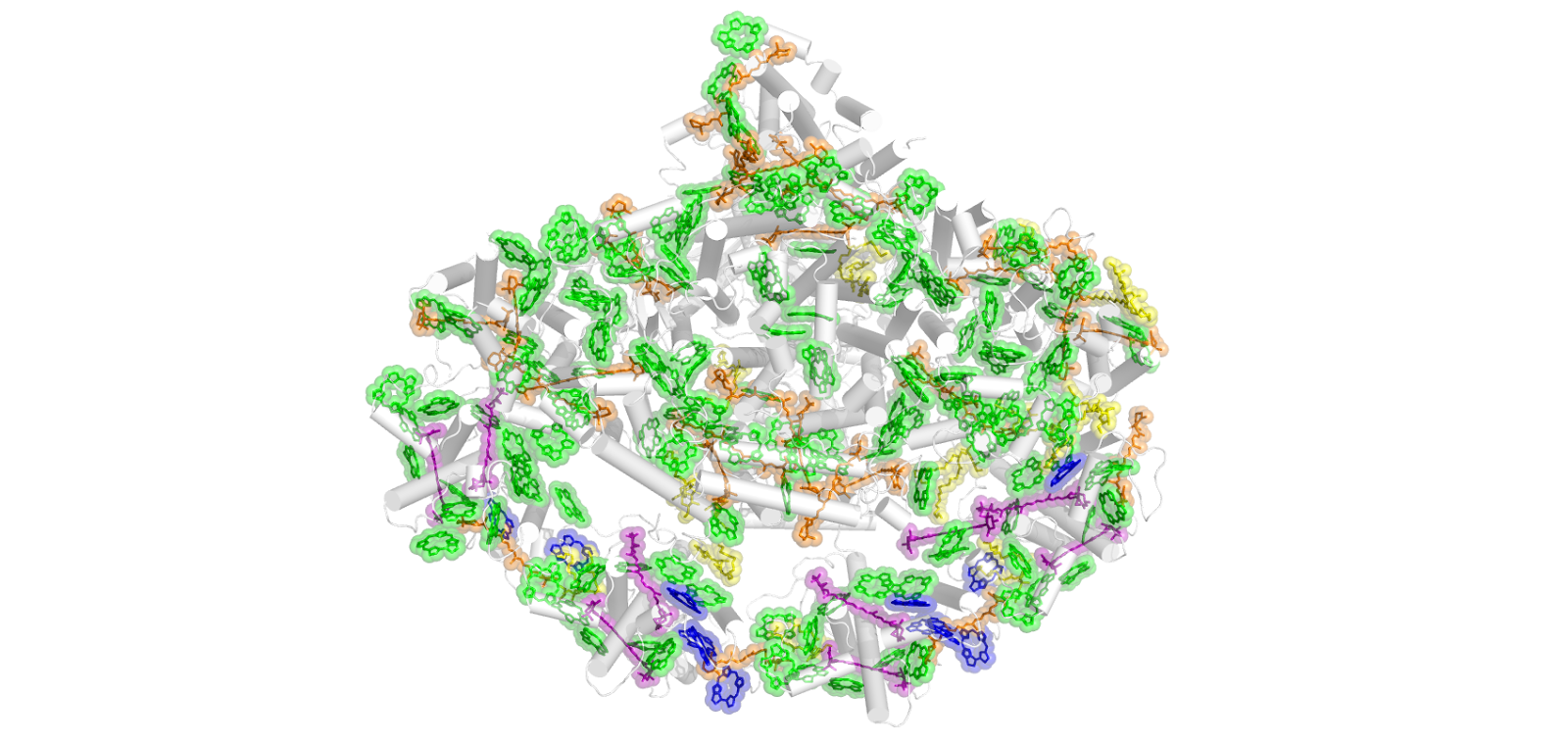

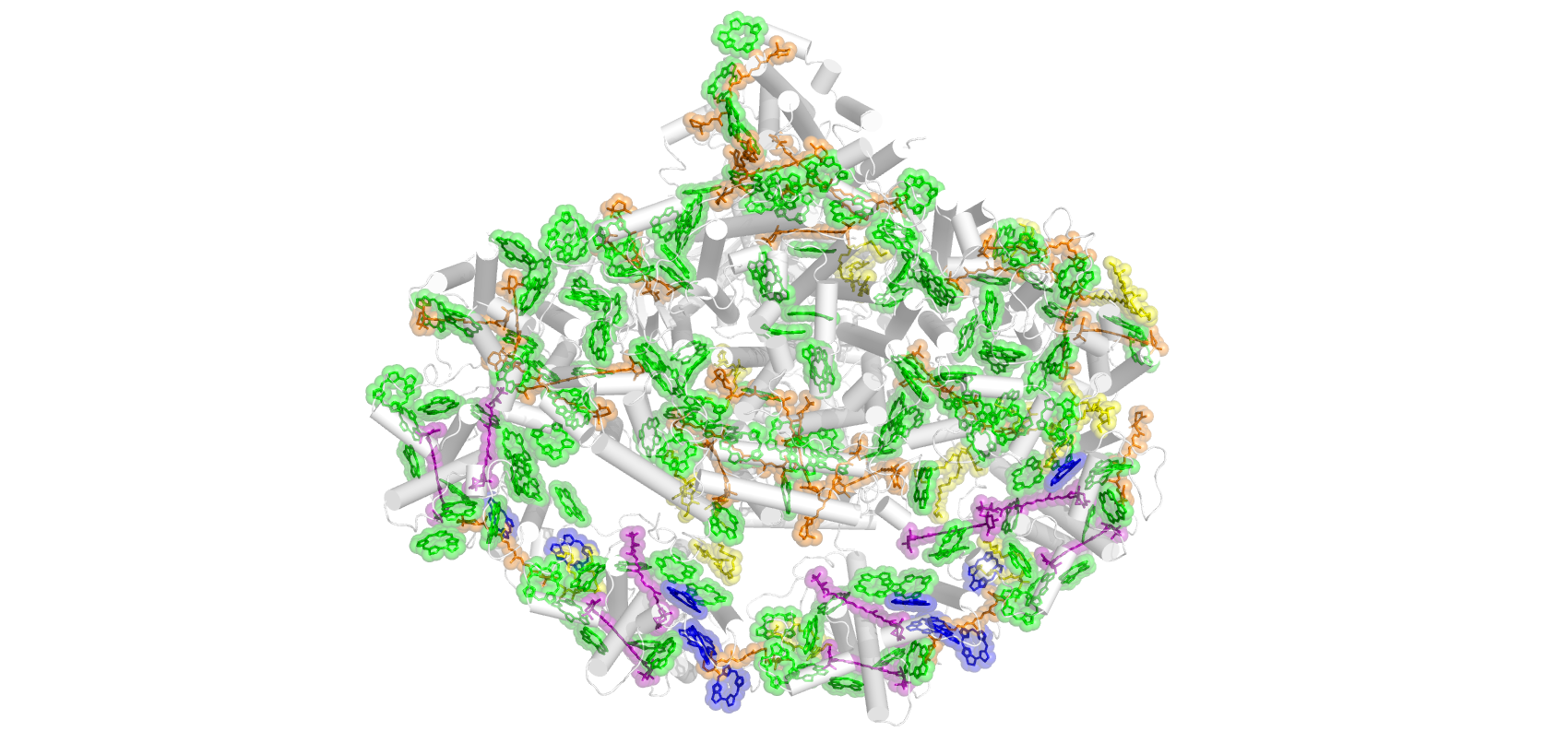

**Figure S8**: Positions of ligands in the PSI-LHCI WT of *A.thaliana* (PDB 9GBI): chl *a* (green), chl *b* (blue), Xanthophylls (purple), β-carotenes (orange), and lipids (yellow).

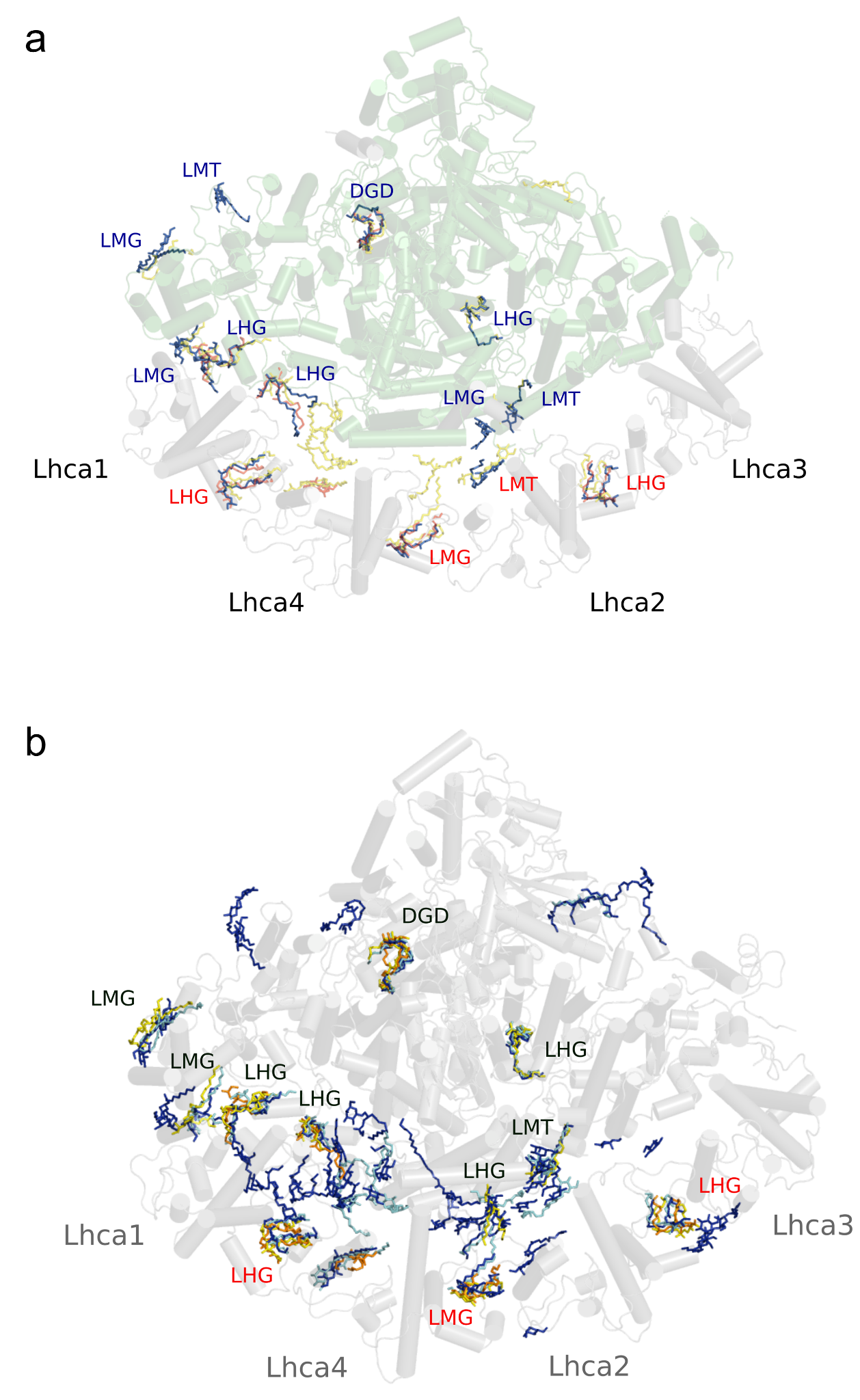

**Figure S9:** **a)** Lipids found in the structures of AtPSI-WT (PDB 9GBI, blue), PDB 8JZB (red), and PDB 7DKZ (yellow). DGD: digalactosyl-diacyl glycerol (DGDG), LMG: 1,2-distearoyl-monogalactosyl-digliceride, LHG: 11,2-dipalmitoyl-phosphatidyl-glycerol, LMT: dodecyl-β-D-maltoside. **b)** Lipids found in the structures of AtPSI-603-NH (PDB 9GC2, yellow), PDB 8JZB (orange), PDB 7DKZ (cyan), and PDB 5L8R (blue). DGD: digalactosyl-diacyl glycerol (DGDG), LMG: 1,2-distearoyl-monogalactosyl-digliceride, LHG: 11,2-dipalmitoyl-phosphatidyl-glycerol, LMT: dodecyl-β-D-maltoside.

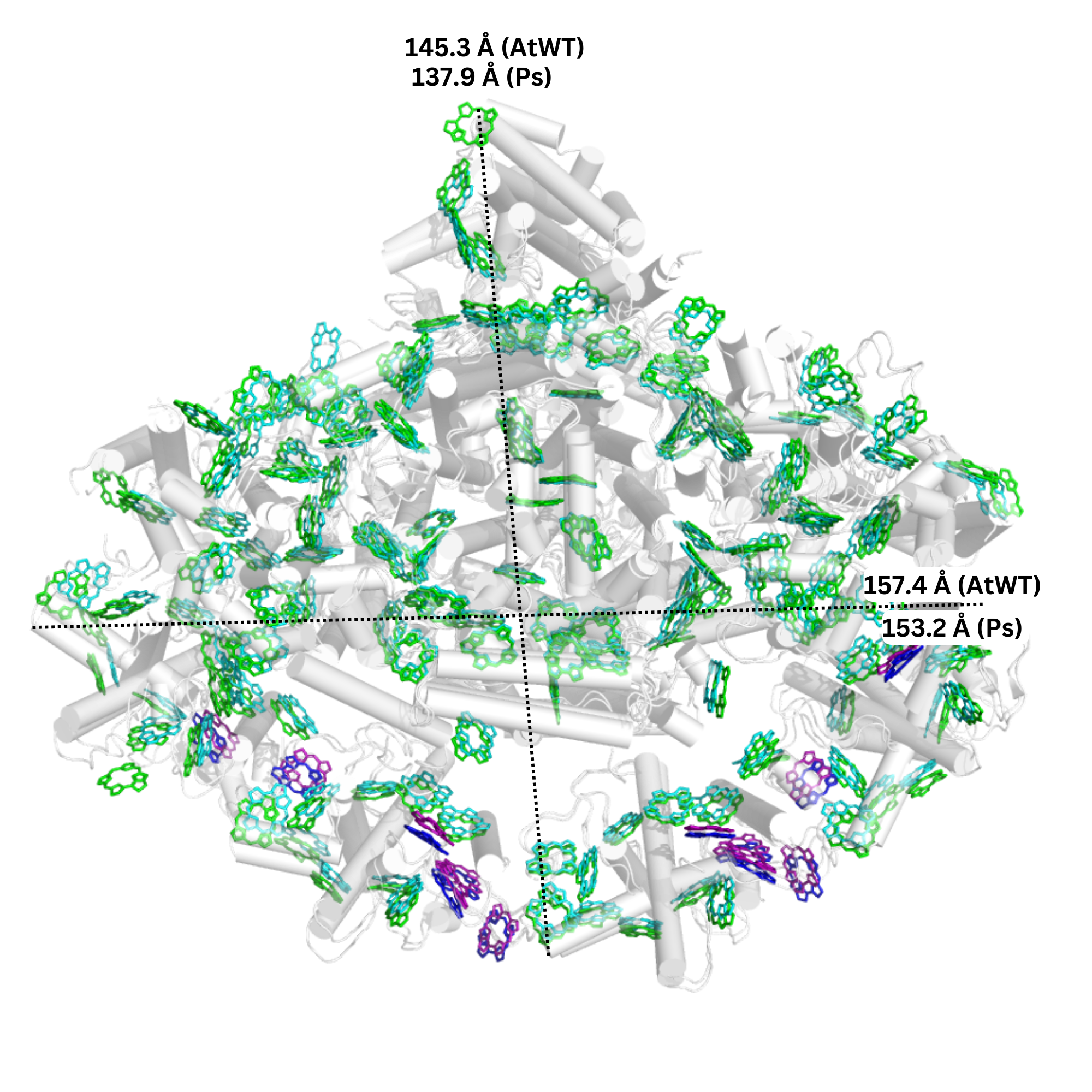

**Figure S10**: Superposition of the PSI WT from *A. thaliana* (PDB 9GBI, white) and from *P. sativum* (PDB 7DKZ). chl *a* from *A. thaliana* and *P. sativum* are colored green and cyan, respectively. chl *b* from *A. thaliana* and *P. sativum* are colored in blue and purple, respectively. The diameter lengths are reported in Å.

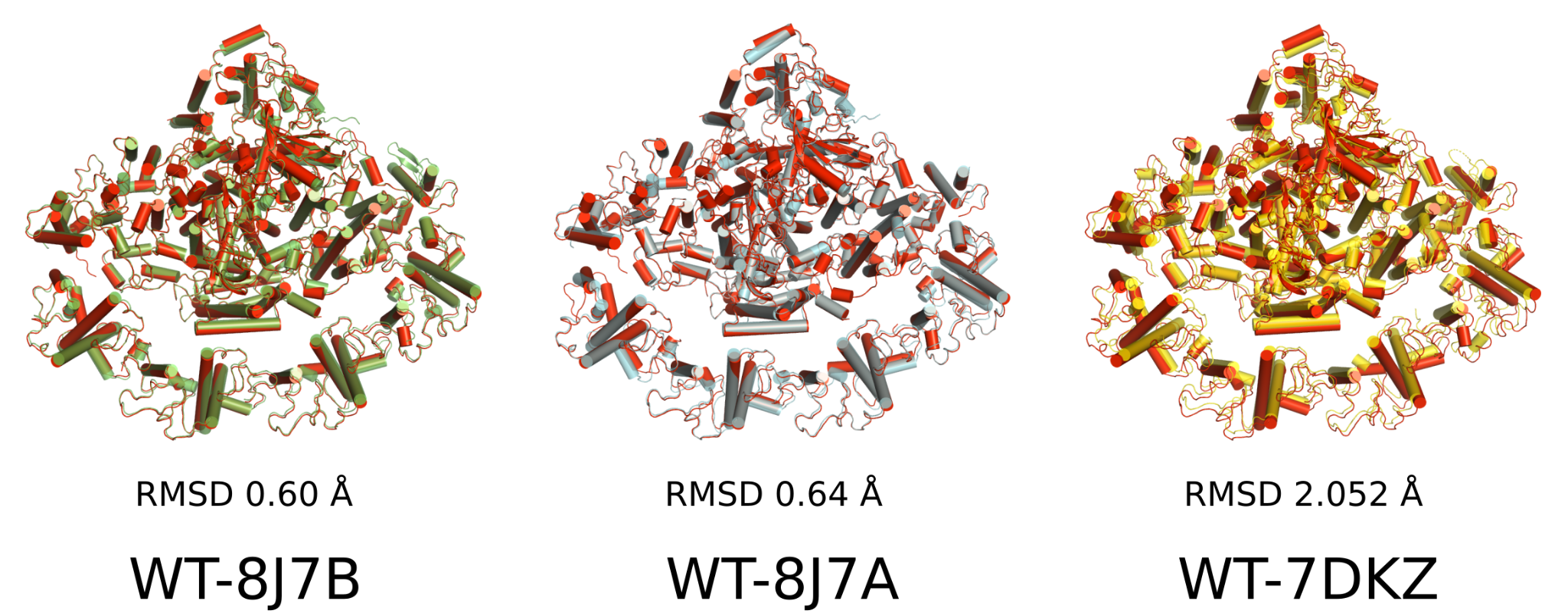

**Figure S11:** Global superposition RMSD of the AtPSI-WT (PDB 9GBI) structure with PSI-WT structures from PDB 8J7B, 8JZA (both from cryo-em data), and 7DKZ (from crystallographic data).

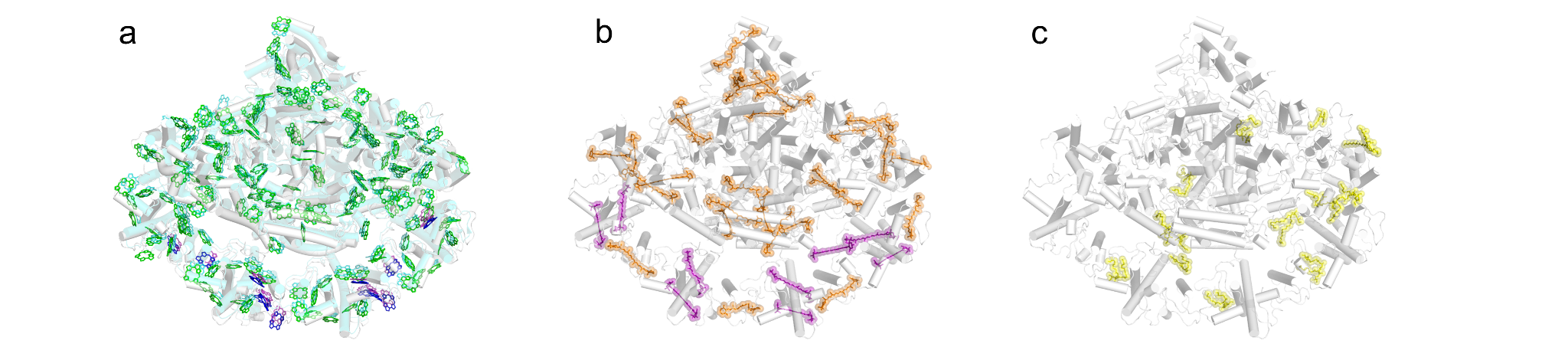

**Figure S12**: **a)** Positions of chl *a* (green) and chl *b* (blue) in the PSI-LHCI WT of *A.thaliana*. **b)** Positions of Xanthophylls (purple) and β-carotenes (orange) in the PSI-LHCI WT of A. thaliana. **c)** Positions of lipids (yellow) in the PSI-LHCI WT of *A. thaliana* (PDB 9GBI).

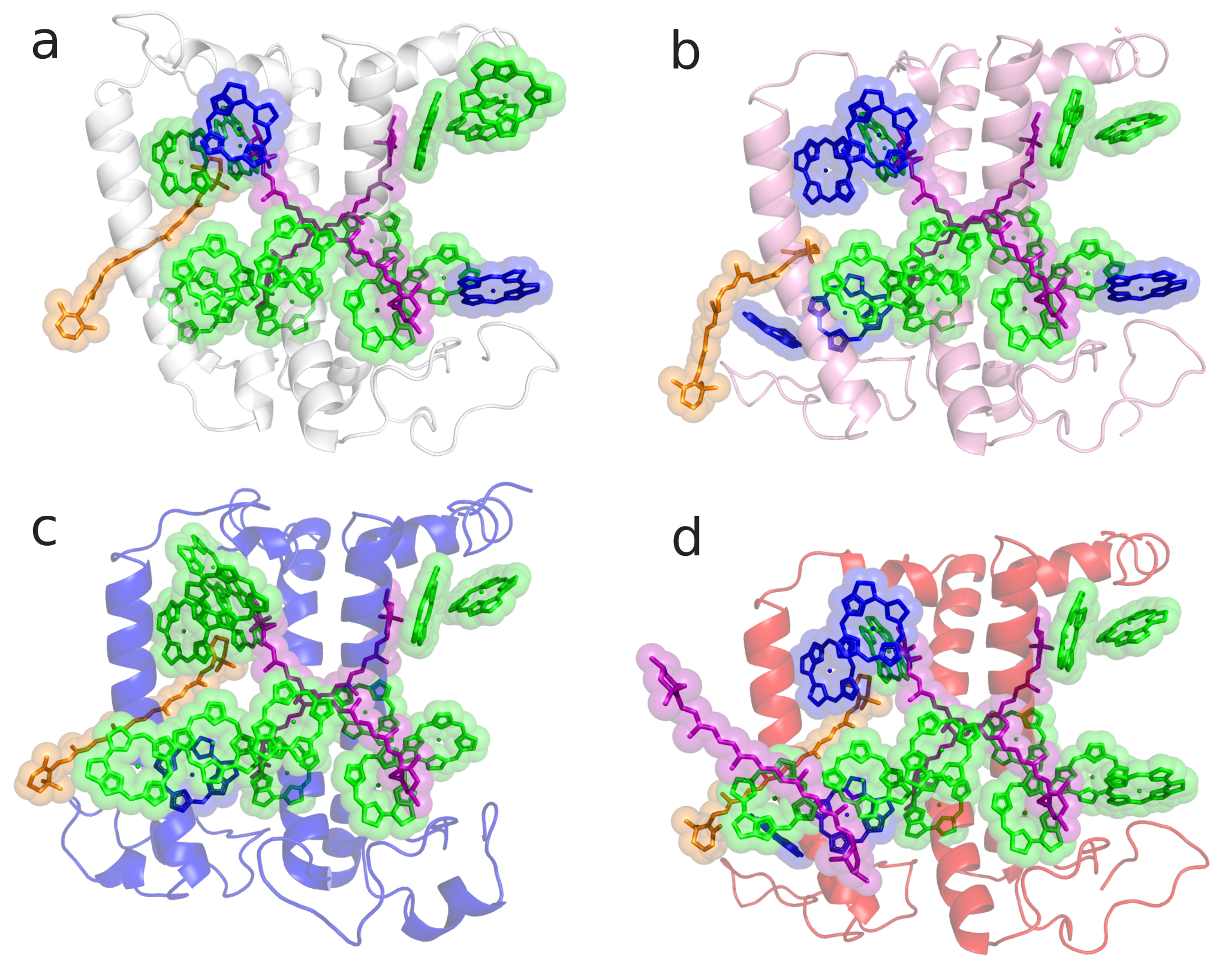

**Figure S13:** Pigment content of the LHCI antenna subunits of *A. thaliana* WT (PDB 9GBI): a) Lhca1, b) Lhca2, c) Lhca3, d) Lhca4. chl *a*, chl *b*, xanthophylls, and β-carotenes are colored green, blue, purple, and orange, respectively.

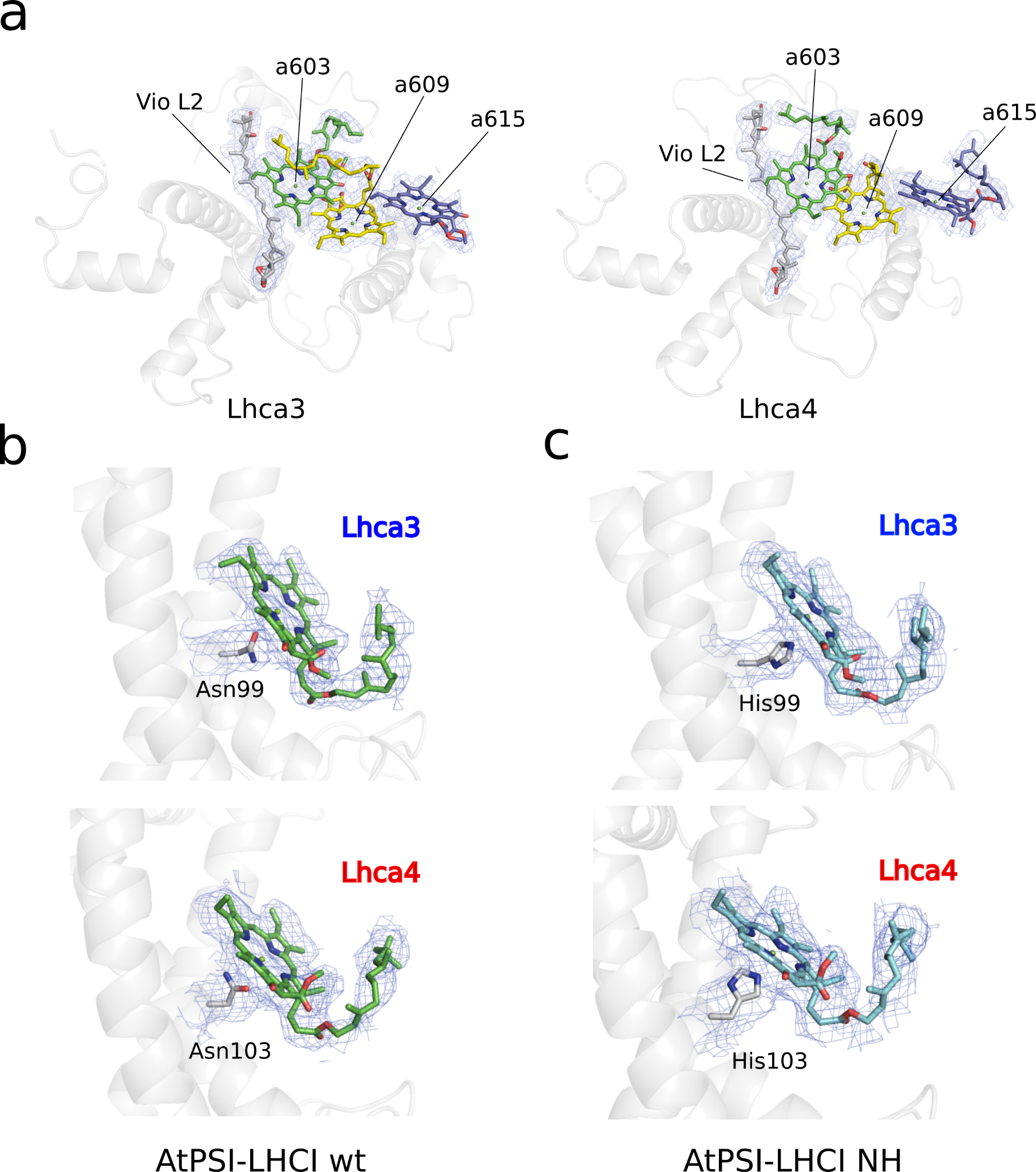

**Figure S14:** a) Atomic models of the “red cluster” pigments in Lhca3 (left) and Lhca4 (right) surrounded by cryo-EM maps. b-c) The position of chl *a*603 and its axial ligand (stick models) in Lhca3 and Lhca4 (WT and *a*603-NH) surrounded by cryo-em maps.

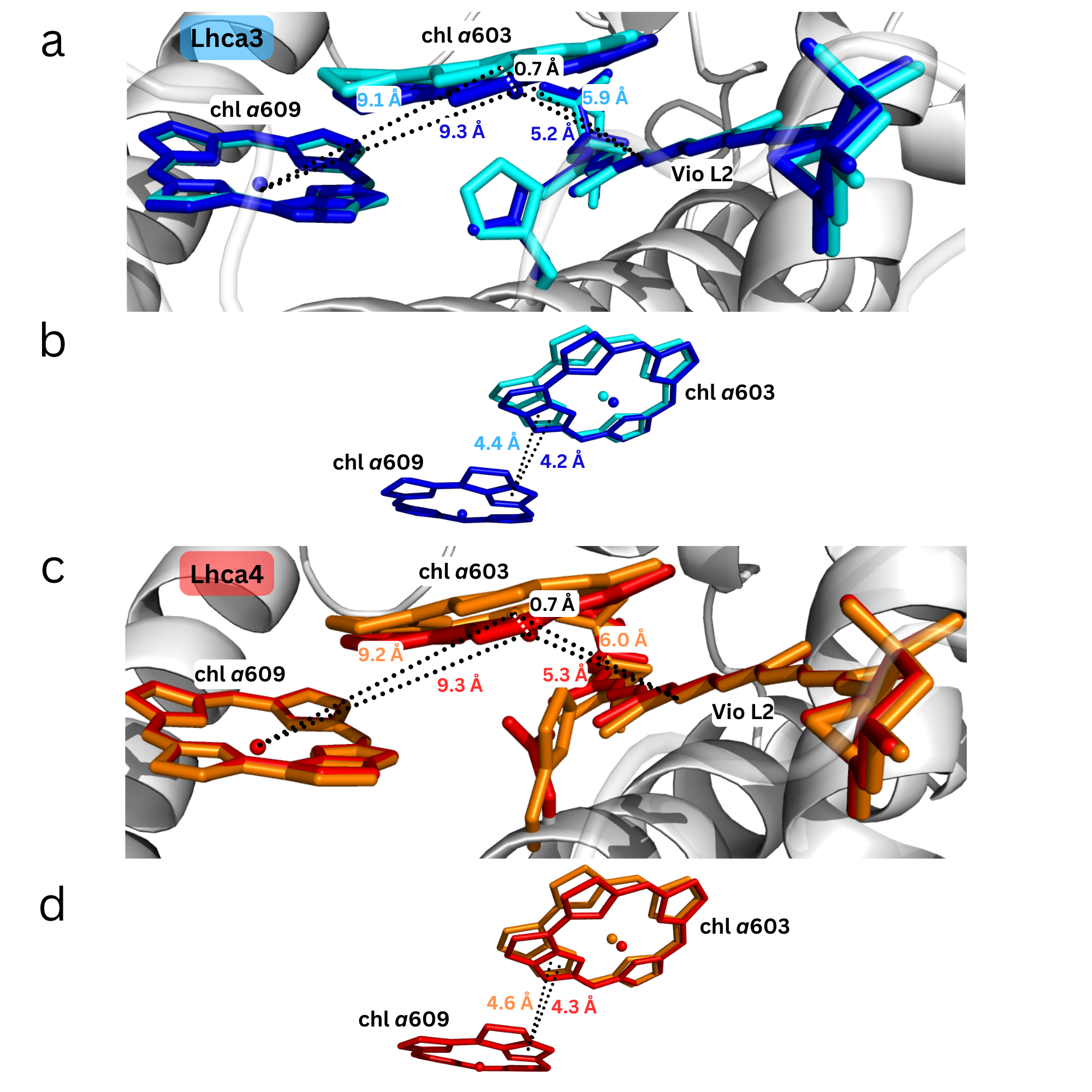

**Figure S15:** a, c) Distances between chl *a*603, chl *a*609, and Violaxanthin L2 and b, d) distances between chlorin C rings of chl *a*603 and chl *a*609 in Lhca3 and Lhca4 subunits of *A. thaliana*. WT (PDB 9GBI) is colored blue and red, and *a*603-NH (PDB 9GC2) is colored cyan and orange.

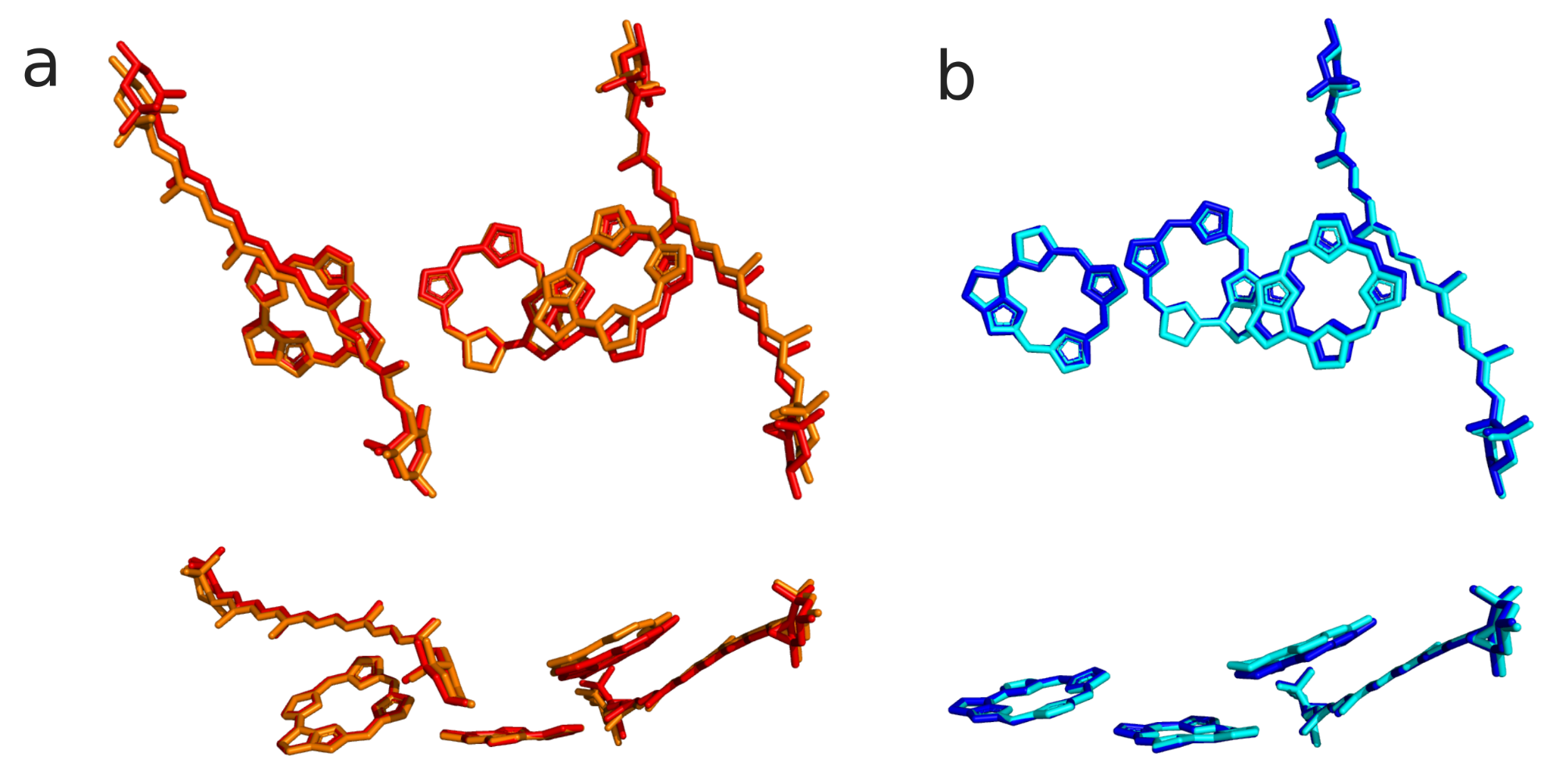

**Figure S16:** Top and side view of superimposed structures of chl *a*603, chl *a*609, chl *a*615, Violaxanthin L2 and Lutein from a) Lhca4 WT and *a603-NH* (red and orange, respectively) and b) Lhca3 WT and *a*603-NH (blue and cyan, respectively). Pigments were superimposed on chl *a*609.

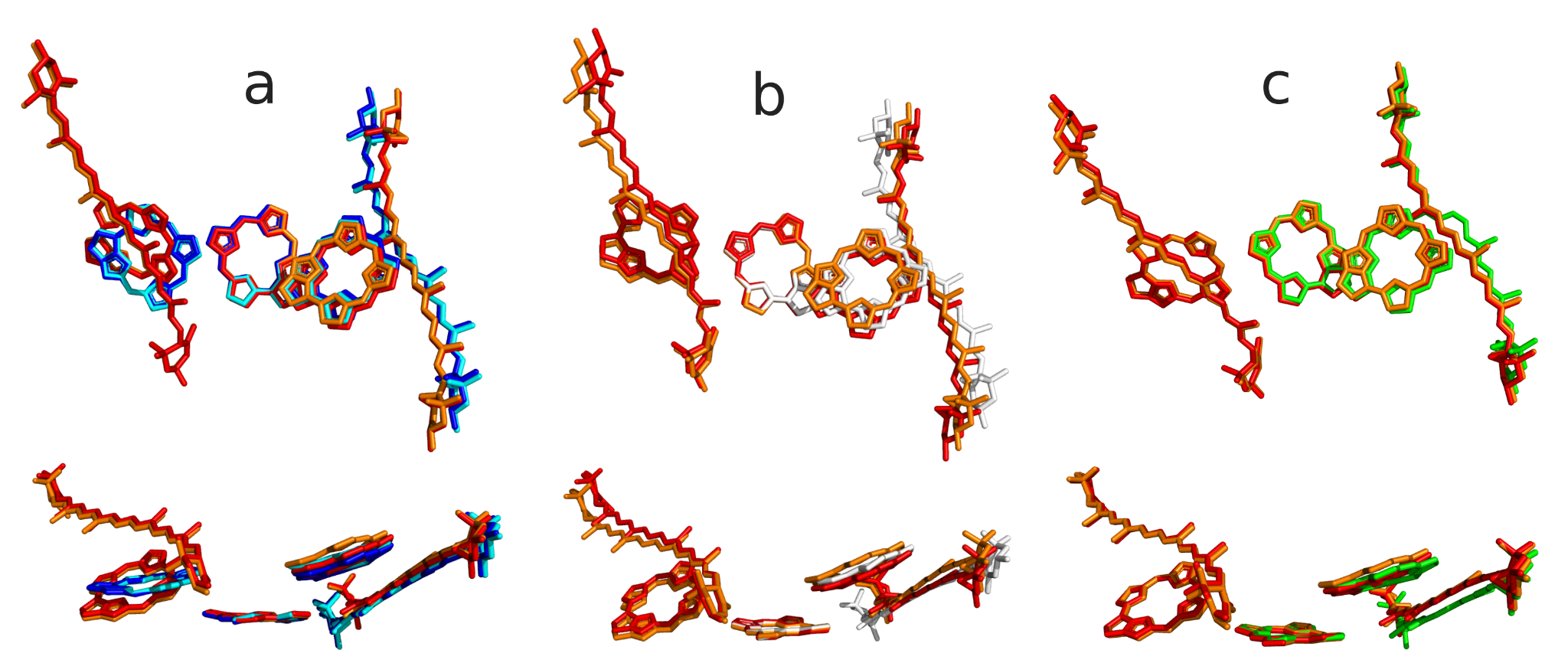

**Figure S17:** Top and side view of superimposed structures of chl *a*603, chl *a*609, chl *a*615, Violaxanthin L2 and Lutein from a) Lhca4 WT and *a603-NH* (red and orange, respectively) with Lhca3 WT and *a603-NH* (blue and cyan, respectively), b) Lhca4 WT and *a603-NH* (red and orange, respectively) with Lhca1 WT (white), and c) Lhca4 WT and *a603-NH* (red and orange, respectively) with Lhca2 WT (green). Pigments were superimposed on chl *a*609.

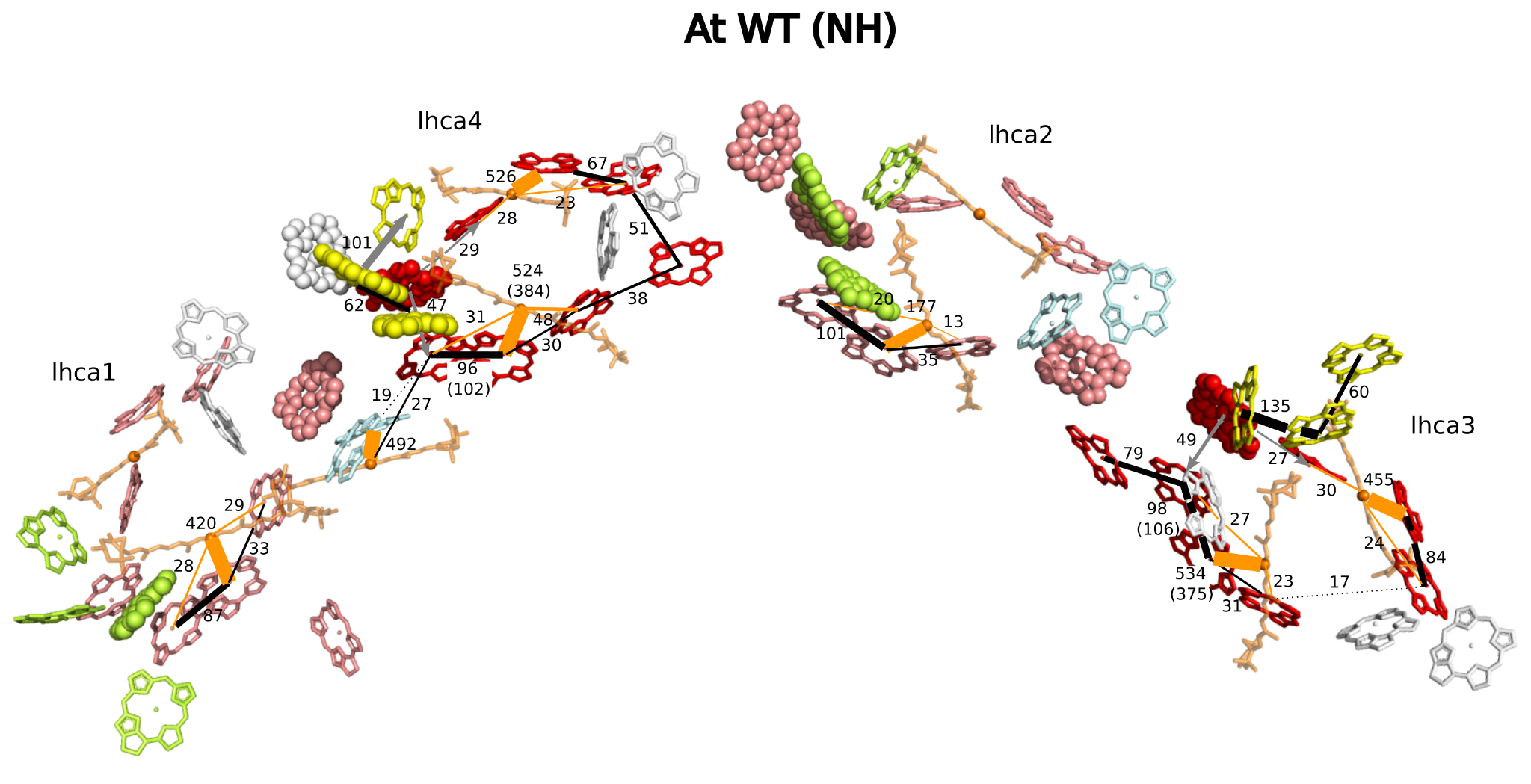

**Figure S18:** Excitonic coupling absolute values between pigments in LHCI WT and *a*603-NH (values in parenthesis). All values are in cm^-1^. Couplings involving xanthophylls and carotenoids are colored orange. The thickness of the line is proportional to the value of the coupling. Couplings with absolute values higher or lower than 20 cm^-1^ are indicated with a solid or dotted line, respectively. Chlorophylls *b* are represented as spheres. Chlorophylls are grouped and colored by clusters wherever the couplings between them are lower than 20 cm^-1^.

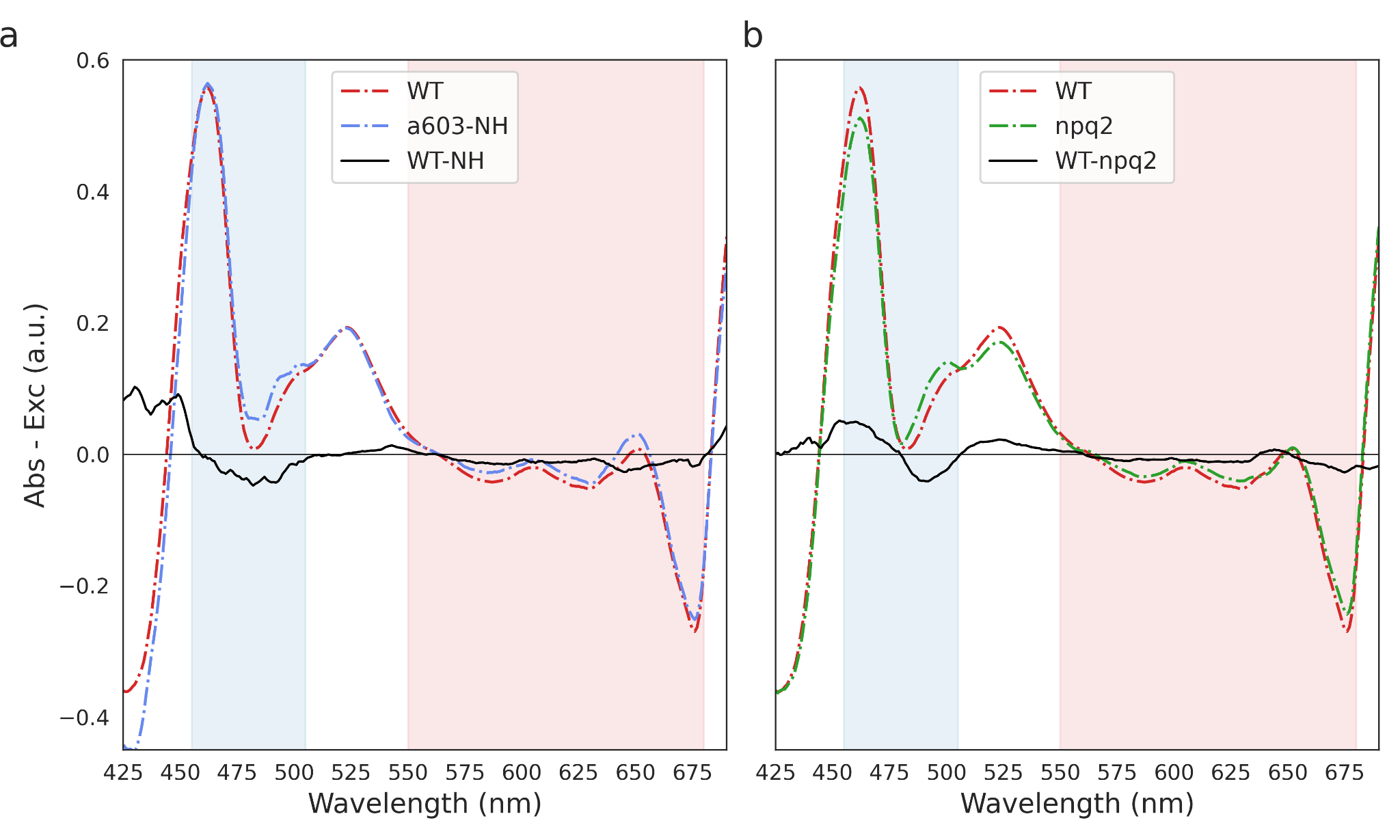

**Figure S19:** Difference spectrum between RT absorption and 77 K excitation spectra in the 425-690 nm region of PSI-LHCI from *A. thaliana* WT and a603-NH (panel a) and *npq2* (panel b) mutant strains. Shaded areas correspond to spectral regions with a prevalence of cars and chls absorption, colored in blue and red, respectively.

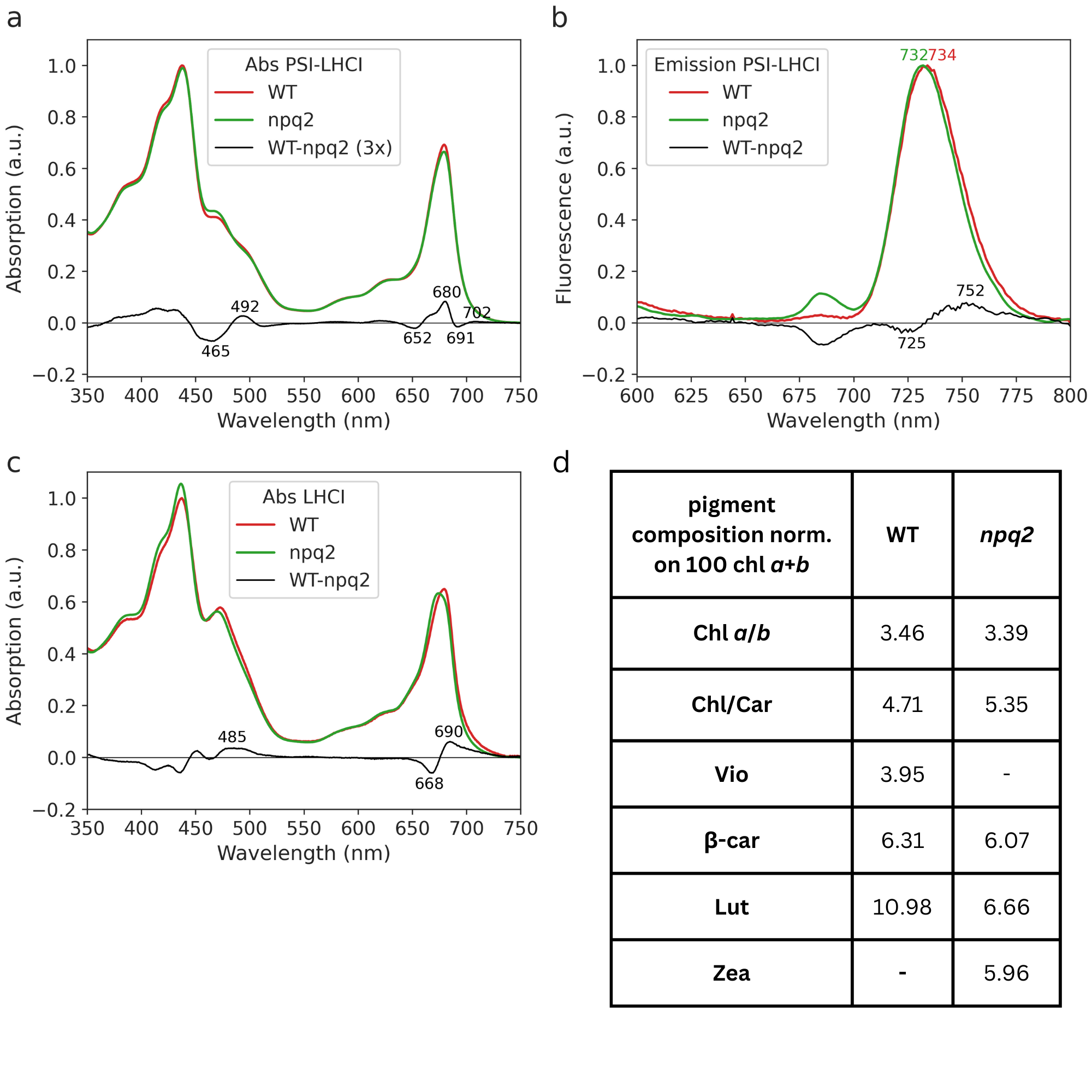

**Figure S20:** (a, c) RT absorption spectra of PSI-LHCI and LHCI from *A. thaliana* WT and *npq2* lines, normalized on the total absorption area. (b) Fluorescence emission spectra of PSI-LHCI from *A. thaliana* WT and *npq2* strains, normalized on the reddest peak. Samples were excited at 440 nm, and emission spectra were recorded from 600 to 800 nm. The difference spectra are shown as black lines in the corresponding plots. The values of the absorption difference spectrum have been magnified by a factor of 3 (panel a) for better visualization. Key wavelengths corresponding to absorption/emission peaks are indicated in nm above the respective peaks. Absorption spectra were normalized on the total absorption area, while the emission spectra were normalized on the maximum. d) Pigment composition of LHCA complexes purified from WT and *npq2*. Quantification was carried out by fitting acetonic spectra and HPLC separation. Car content was normalized on 100 chls *a*+*b*.

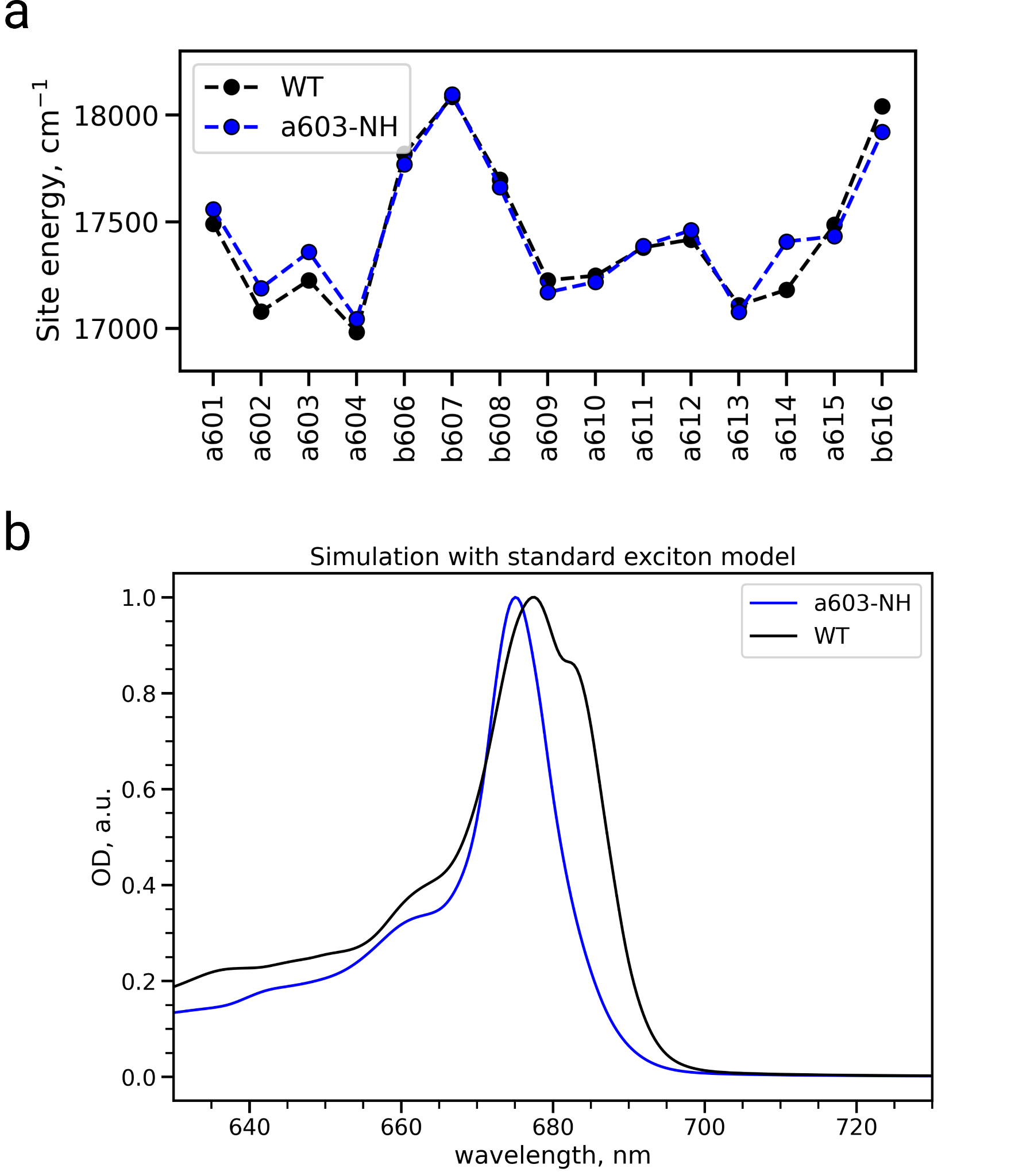

**Figure S21:** (a) Site energies of Lhca4 chls for WT and *a*603-NH mutant. (b) Absorption spectra of WT and mutant simulated using the simple exciton model (local excitations only, without including the CT states).

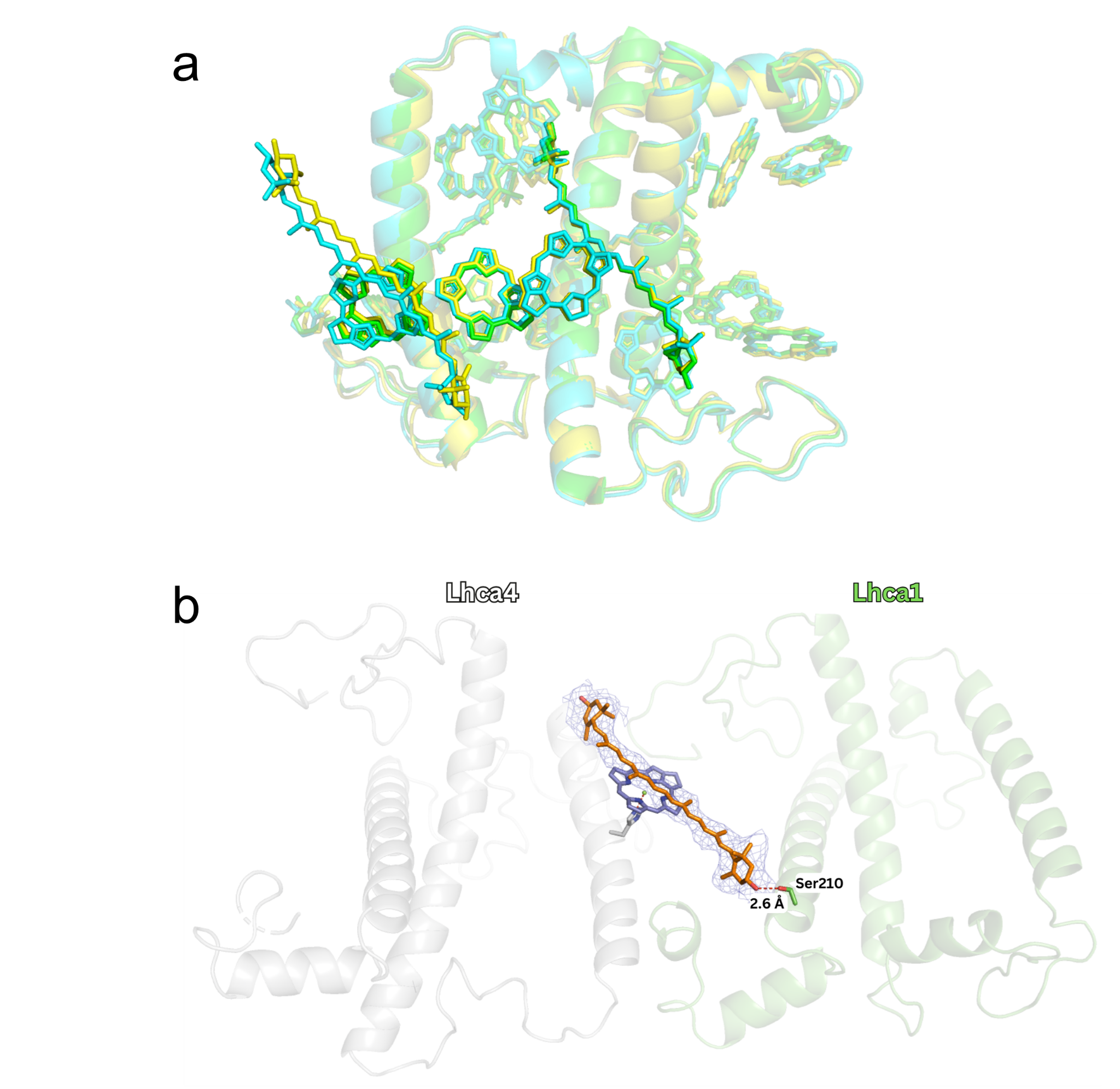

**Figure S22:** **a)** Structural superposition of Lhca4 WT from *A. thaliana* (cyan, PDB 9GBI), *P. sativum* (green, PDB 5L8R), *Z. mays* (yellow, PDB 5ZJI). The “red cluster” formed by chl *a*603, chl *a*609, and chl *a*615 is shown from right to left. The extra lutein is found in the vicinity of chl *a*615 only in *A. thaliana* and *Z. mays*. **b)** Cryo-em map for Lut over chl a615 in Lhca4. Focus on the hydrogen bond of the Lut hydroxyl with Ser210 from Lhca1.

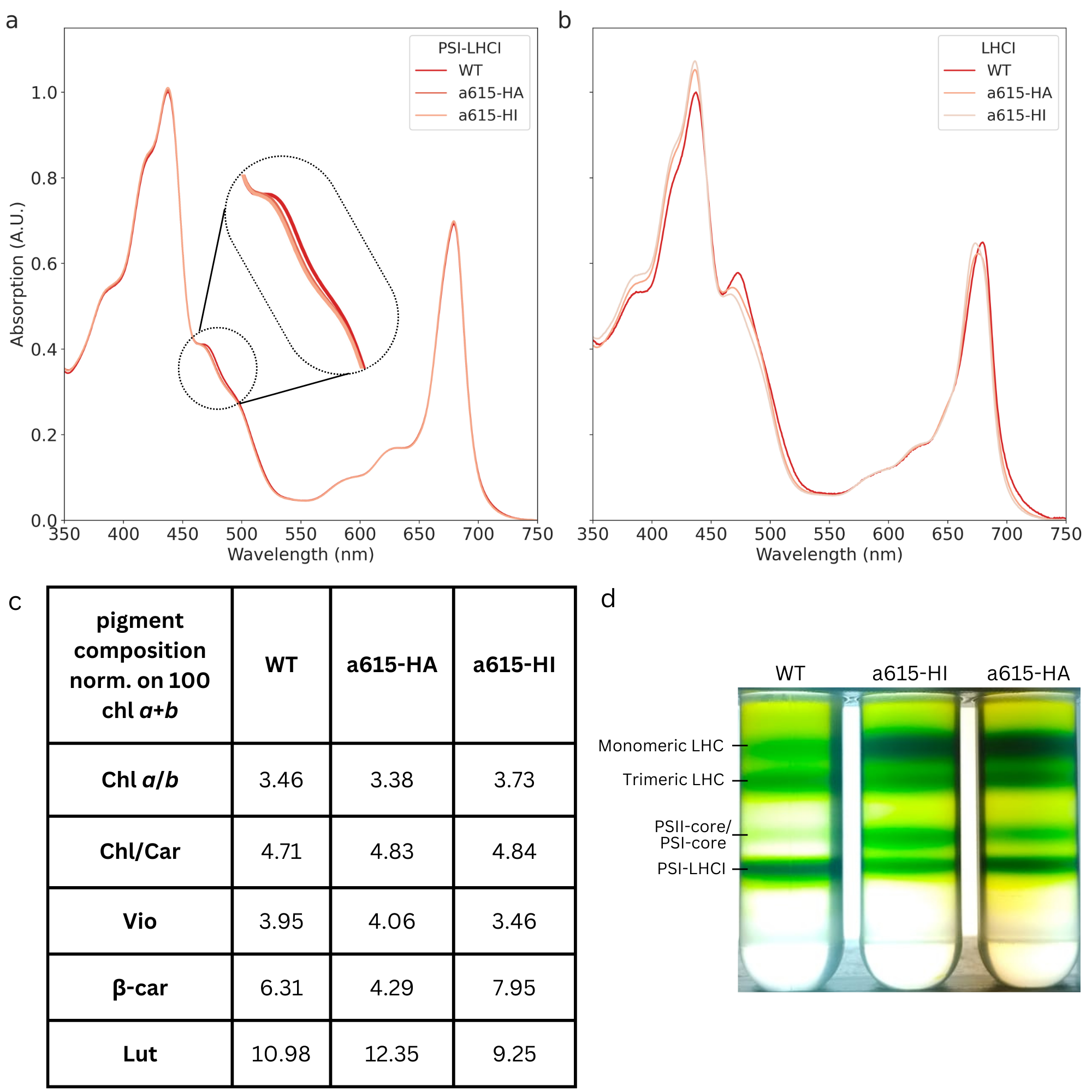

**Figure S23:** Sucrose gradient fractionation of thylakoid membranes of WT and *a*615 mutant lines solubilized with 1% β-DM.

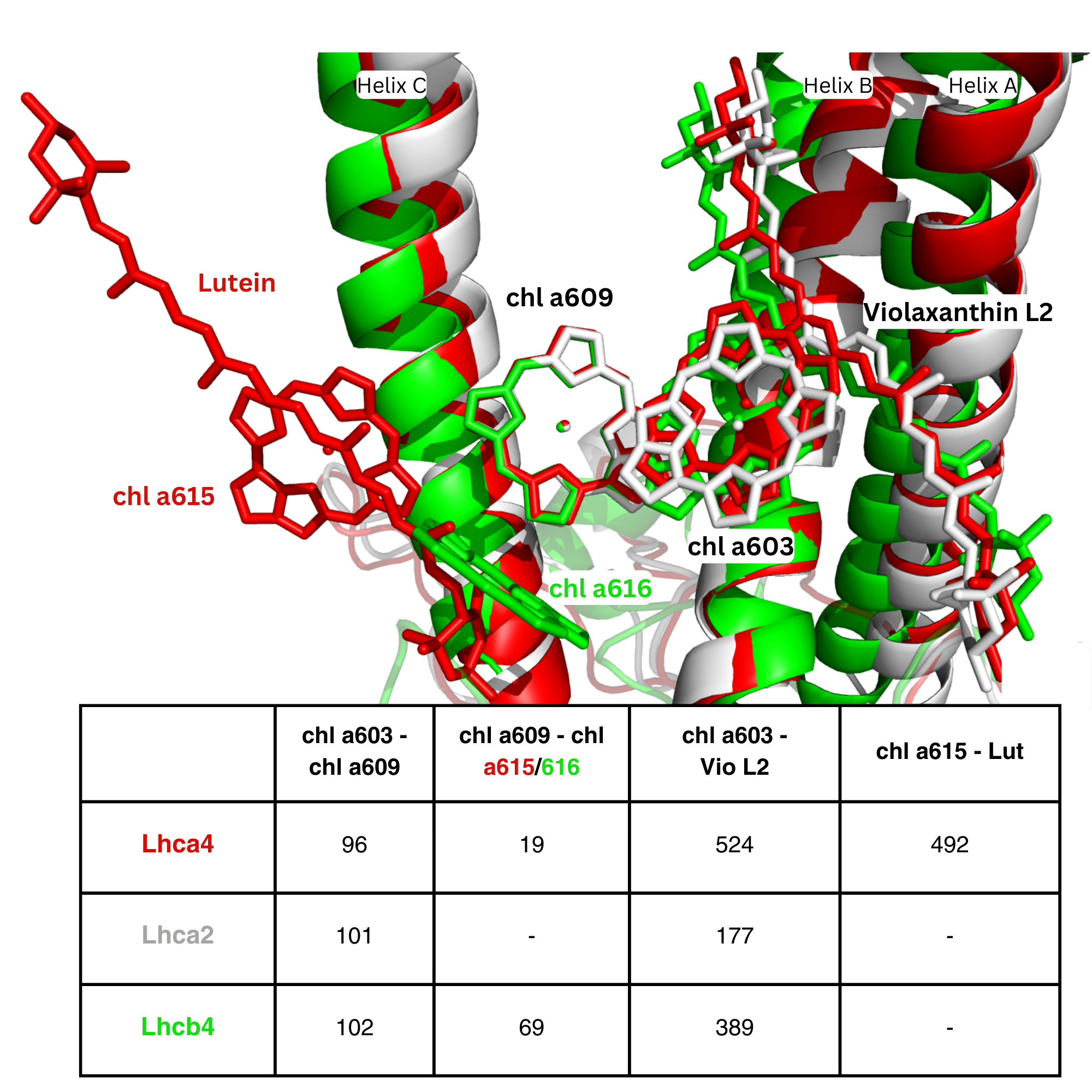

**Figure S24:** Superposition of Lhca4, Lhca2 from *A. thaliana* WT (PDB 9GBI) and Lhcb4 from *P. sativum* (PDB 5XNL) and excitonic couplings between pigments pairs in the red cluster.

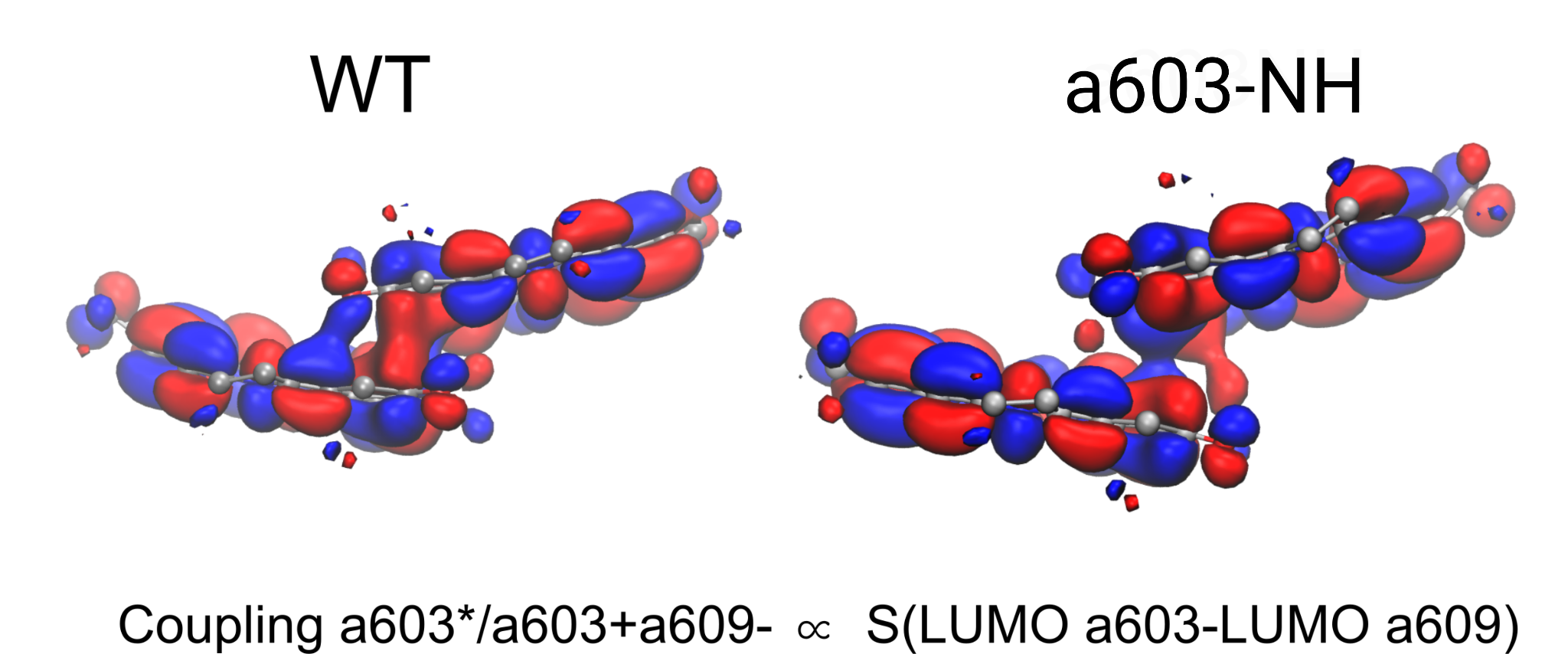

**Figure S25:** Visual representation of the overlap between LUMO orbitals of chls *a*603 and *a*609 as computed for the WT and *a*603-NH structures.

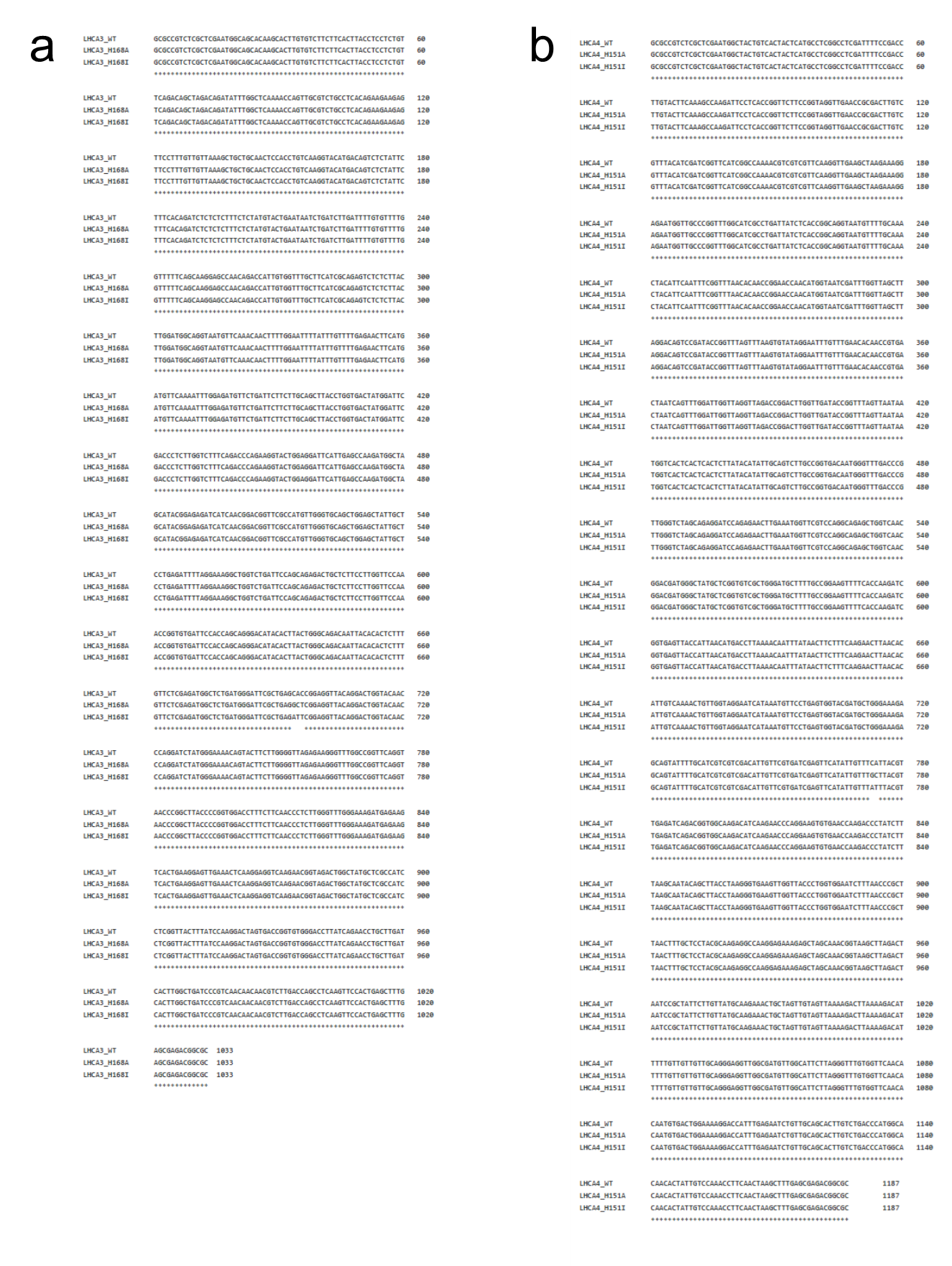

**Figure S26:** **a**. Sequence alignment of the sequences (exons and introns) of the synthetic genes encoding for *Lhca3_WT* and *Lhca3_a615-H168A* and *Lhca3_a615-H168I*. **b**. Sequence alignment of coding sequences and introns of the synthetic genes encoding for *Lhca4_WT* and *Lhca4_a615-H151A* and *Lhca4_a615-H151I*.

| **Subunit** | **Traced**  **residues** | **Chlorophylls *a*** | **Chlorophylls *b*** | **carotenoids** | **Lipids** | **Others** |
| --- | --- | --- | --- | --- | --- | --- |
| PsaA | 742 | 41 CLA |  | 7 BCR | 1 LHG,  1 LMT | 1 PQN,  1 SF4 |
| PsaB | 733 | 41 CLA |  | 6 BCR | 1 LHG,  1 LMG,  1 DGD | 1 PQN |
| PsaC | 80 | - | - | - | - | 2 SF4 |
| PsaD | 141 | - | - | - | - | - |
| PsaE | 63 | - | - | - | - | - |
| PsaF | 152 | 3 CLA | - | 3 BCR | 1 LHG | - |
| PsaG | 94 | 3 CLA | - | 1 BCR | 1 LHG,  1 LMT | - |
| PsaH | 89 | 1 CLA | - | - | - | - |
| PsaI | 30 | - | - | 1 BCR | - | - |
| PsaJ | 39 | 2 CLA | - | 2 BCR | - | - |
| PsaK | 50 | 2 CLA | - | - | - | - |
| PsaL | 144 | 3 CLA | - | 3 BCR | - |  |
| PsaN | 48 | - | - | - | - | - |
| Lhca1 | 194 | 12 CLA | 2 CHL | 1 XAT,  1 LUT,  1 BCR | 1 LHG | - |
| Lhca2 | 202 | 9 CLA | 5 CHL | 1 XAT,  1 LUT,  1 BCR | 1 LHG | - |
| Lhca3 | 216 | 13 CLA | 1 CHL | 1 XAT,  1 LUT,  1 BCR | - | - |
| Lhca4 | 195 | 11 CLA | 4 CHL | 1 XAT,  2 LUT,  1 BCR | 1 LMG | - |

**Table S1**: AtPSI-603-NH structural model. CLA, chl *a*; CHL, chl *b*; BCR, β-carotene; LUT, lutein; XAT, violaxanthin; DGD, digalactosyl-diacyl glycerol (DGDG); LMG, 1,2-distearoyl-monogalactosyl-digliceride; LHG, 11,2-dipalmitoyl-phosphatidyl-glycerol; LMT, dodecyl-β-D-maltoside; PQN, phylloquinone; SF4, Fe4-S4 cluster.

| **Primer Name** | **Sequence** | **Purpose** |
| --- | --- | --- |
| **Lhca3_N103H_FW** | CATACGGAGAGATCATCCATGGACGGTTCGCCATG | Site-directed mutagenesis of Lhca3 N103H |
| **Lhca3_N103H_RV** | CATGGCGAACCGTCCATGGATGATCTCTCCGTATG | Site-directed mutagenesis of Lhca3 N103H |
| **Lhca4_N99H_FW** | GGCAGAGCTGGTCCATGGACGATGGGCTATG | Site-directed mutagenesis of Lhca4 N99H |
| **Lhca4_N99H_RV** | CATAGCCCATCGTCCATGGACCAGCTCTGCC | Site-directed mutagenesis of Lhca4 N99H |
| **Lhca3_seq_FW** | TCTTCTACCCTAACATAGCCCTTGC | Sequencing of Lhca3 genomic sequence |
| **Lhca3_seq_RV** | AGAAGCCAATGGTAGCGAAAGATGG | Sequencing of Lhca3 genomic sequence |
| **Lhca4_seq_FW** | CCAATCCATTCTTCTTCAAGTGCC | Sequencing of Lhca4 genomic sequence |
| **Lhca4_seq_RV** | TCACAGACAGACATGAAAGTGATGG | Sequencing of Lhca4 genomic sequence |

**Table S2:** List of the primers used to obtain and characterize the *a603-NH* mutant lines.

|  | **AtPSI-WT** | **AtPSI-603-NH** |
| --- | --- | --- |
| **Data collection and**  **processing** |  |  |
| Magnification | 120,000X | 120,000X |
| Voltage (kV) | 200 | 200 |
| Total electron dose (e-/Å) | 40 | 40 |
| Nominal defocus range（µm） | 0.8 - 2.4 | 0.8 - 2.4 |
| Pixel size（Å） | 0.889 | 0.889 |
| Symmetry | C1 | C1 |
| Initial particle images (no.) | 575,271 | 115,589 |
| Final particle images (no.) | 54,155 | 36,596 |
| Map resolution (Å) | 3.13 | 3.29 |
| FSC threshold | 0.143 | 0.143 |
| **Refinement** |  |  |
| Initial model used (PDB code) | 7DKZ | 9GBI |
| Map sharpening B factor (Å^2^ ) | 83.7 | 91.8 |
| Model composition |  |  |
| Protein residues | 3230 | 3190 |
| Ligands | 210 | 206 |
| B factor (Å^2^) |  |  |
| Protein | 51.02 | 70.22 |
| Ligands | 49.35 | 68.56 |
| R.m.s. deviations |  |  |
| Bond lengths (Å) | 0.002 | 0.003 |
| Bond angles (°) | 0.518 | 0.551 |
| **Validation** |  |  |
| MolProbity score | 1.51 | 1.48 |
| Clash score | 7.19 | 7.53 |
| Rotamers outliers (%) | 0.34 | 0.27 |
| Ramachandran plot |  |  |
| Favored (%) | 97.4 | 97.7 |
| Allowed (%) | 2.6 | 2.3 |
| Disallowed (%) | 0 | 0 |
| CC (model vs data) |  |  |
| Mask | 0.88 | 0.88 |
| Box | 0.69 | 0.72 |

**Table S3:** Cryo-EM data collection, refinement, and validation statistics.
